# ASCL1 interacts with the mSWI/SNF at distal regulatory elements to regulate neural differentiation

**DOI:** 10.1101/2022.10.09.510609

**Authors:** Oana Păun, Yu Xuan Tan, Harshil Patel, Stephanie Strohbuecker, Avinash Ghanate, Clementina Cobolli-Gigli, Miriam Llorian Sopena, Lina Gerontogianni, Robert Goldstone, Siew-Lan Ang, François Guillemot, Cristina Dias

**Affiliations:** Neural Stem Cell Biology Laboratory, The Francis Crick Institute, 1 Midland Road, London NW1 1AT, UK; Bioinformatics & Biostatistics Science and Technology Platform, The Francis Crick Institute, 1 Midland Road, London NW1 1AT, UK; Medical and Molecular Genetics, School of Basic and Medical Biosciences, Faculty of Life Sciences & Medicine, King’s College London, London SE1 9RT, UK

**Keywords:** Neural stem cell, iPSC-derived culture, neurogenesis, ASCL1, mSWI/SNF, chromatin regulation, pioneer transcription factor

## Abstract

Pioneer transcription factors are thought to play pivotal roles in developmental processes by binding nucleosomal DNA to activate gene expression. The role of neurogenic pioneer factor ASCL1 in shaping chromatin landscape in human neurogenesis remains unclear. Here we show that ASCL1 acts as a pioneer transcription factor in a transient population of progenitors. Using an *in vitro* ASCL1 knockout model we show it drives progenitor differentiation by cis-regulation both as a classical pioneer factor and as a non-pioneer remodeler, where ASCL1 binds permissive chromatin to induce chromatin conformation changes. We find ASCL1 directly interacts with mammalian BAF SWI/SNF chromatin remodeling complexes, essential for neurogenesis and involved in multiple neurodevelopmental disorders. ASCL1 acts as a non-pioneer chromatin remodeler to regulate gene expression at a subset of loci, requiring mBAF SWI/SNF’s ATPase activity for cis-regulation of gene expression. Our findings demonstrate that ASCL1 is a key chromatin remodeler in human neurogenesis, uncovering an alternative mechanism of remodeling function dependent on partner ATPase activity.

## Introduction

Neuronal differentiation of stem and progenitor cells in the developing mammalian brain is a highly regulated process that results in the generation of neurons of distinct identities in appropriate numbers and at defined locations. A group of transcription factors (TFs) of the basic helix-loop-helix (bHLH) class, called proneural proteins have a prominent role in the regulation of neurogenesis (Bertrand et al., 2002; Huang et al., 2014). In particular, the proneural factor ASCL1 has been shown to regulate multiple steps of neurogenesis, including the proliferation and neuronal fate commitment of multipotent progenitors, and the differentiation and migration of postmitotic neurons (Borromeo et al., 2014; Castro et al., 2011; Nieto et al., 2001; Pacary et al., 2011; Tomita et al., 2000). ASCL1 function in neurogenesis has been studied mostly in the developing mouse brain (Andersen et al., 2014; Castro et al., 2011; Nieto et al., 2001; Pacary et al., 2011). However, ASCL1 expression has also been reported in progenitor cells in the human embryonic telencephalon (Alzu’bi and Clowry, 2019; Hansen et al., 2010), suggesting that it has retained a function in the regulation of neurogenesis in humans.

Some of the TFs regulating the differentiation of tissues in the embryo control early steps in the development of cell lineages via their pioneer activity (Cirillo et al., 2002; Gao et al., 2019; Iwafuchi-Doi et al., 2016; Mayran et al., 2019; Song et al., 2021; Yu et al., 2021; Zaret and Carroll, 2011). Most TFs interact with their cognate binding sites in the genome only when in an open chromatin configuration, i.e. in a stretch of DNA that is not wrapped around nucleosomes. In contrast, pioneer transcription factors have the capacity to recognize binding sites in regulatory elements embedded in closed chromatin, i.e. in DNA wrapped around nucleosomes, and to open chromatin, thus facilitating the binding of other transcription factors and the transcription of the associated genes (Iwafuchi-Doi and Zaret, 2014; Soufi et al., 2015). A third category has recently been proposed, “non-classical” pioneer TFs, which remodel chromatin but have DNA binding properties constrained by nucleosome position, and require the recruitment of chromatin remodeling complexes to their target sites (Minderjahn et al., 2020).

Proneural proteins, including ASCL1, have been shown to display pioneer activity, recognizing nucleosomal DNA enriched for a short E-box motif (Henke et al., 2009; Soufi et al., 2015). Evidence for the pioneer activity of ASCL1 comes from *in vitro* reconstituted nucleosome binding assays and studies involving overexpression of the protein in cultured cells (Aydin et al., 2019; Chanda et al., 2014; Fernandez Garcia et al., 2019; Heinrich et al., 2010; Park et al., 2017; Raposo et al., 2015; Soufi et al., 2015; Wapinski et al., 2013). In particular, ASCL1 has been shown to recognize its neuronal targets in a closed chromatin state when overexpressed in fibroblasts and thereby initiate neuronal reprogramming (Wapinski et al., 2013). Similarly, ASCL1 binds neuronal genes in a closed configuration when overexpressed in undifferentiated neural stem cells or glioblastoma stem cells (Park et al., 2017; Raposo et al., 2015). However, ASCL1 function in human neurogenesis is currently unknown. Moreover, the pioneer activity of the endogenous ASCL1 protein has not yet been investigated, nor has the mechanism by which local chromatin structure is affected by ASCL1 binding. Different models have been proposed to explain the capacity of pioneer factors to open chromatin, including physical displacement of nucleosomes by the pioneer factor-DNA interaction (Cirillo et al., 2002; Michael et al., 2020), and/or interaction with other transcription factors and proteins with chromatin remodeling capacity (Hu et al., 2011; King and Klose, 2017; Takaku et al., 2016; Theodorou et al., 2013; Wang et al., 2014).

Four families of chromatin remodeling factors have been identified in mammalian cells (Becker and Workman, 2013). Among them, the mammalian mSWI/SNF complexes represent attractive candidates to interact with ASCL1, because of their role in the regulation of neurogenesis, exemplified by their implication in multiple neurodevelopmental disorders (Kosho et al., 2014; Pulice and Kadoch, 2016; Sokpor et al., 2017). While core subunits of BAF complexes are ubiquitously expressed, other subunits are incorporated with developmental stage and cell-type specificity (Mashtalir et al., 2020; Son and Crabtree, 2014). Importantly, changes in the combinatorial assembly of BAF complexes underpin the transition from neural progenitors to neurons *in vivo* and *in vitro* (Lessard et al., 2007; Staahl et al., 2013; Yoo et al., 2009), with specific subunits incorporating in progenitor-specific (npBAF) or neuronal-specific (nBAF) complexes. The developmental overlap of npBAF and proneural transcription factors was recently highlighted by the demonstration that conditional deletion of npBAF subunit ACTL6A in mouse cortex leads to decreased chromatin accessibility at proneural TF binding sites, including ASCL1 (Braun et al., 2021). We therefore hypothesized that the pioneer TF ASCL1 may interact with mBAF SWI/SNF complexes to regulate chromatin accessibility at neurogenic loci to coordinate neurodifferentiation.

In this study, we have investigated the mechanisms by which ASCL1 regulates neurogenesis in a cell culture model of human cortical development. We show that ASCL1 expression defines a cell population of transitional neural progenitors immediately preceding differentiation to neurons. ASCL1 has pioneer activity, binding regulatory elements implicated in neurogenic programs in a closed chromatin configuration and opening chromatin at many of its target sites. It also binds sites with a low degree of accessibility to induce further chromatin changes. Cooperative mSWI/SNF chromatin remodeling ATPase activity is required for ASCL1 activity at a subset of its targets. Together, our findings support a model whereby ASCL1 acts as a pioneer factor in distinct ways during human telencephalic neurogenesis. It acts as a classical pioneer transcription factor binding heterochromatin for regulation of a fraction of its targets, largely without mSWI/SNF interaction. However, it also regulates chromatin accessibility at a large number of loci in the genome via cooperative binding with the mSWI/SNF chromatin remodelers at sites of permissive DNA.

## Results

### ASCL1 is expressed in cells transitioning from dividing progenitors to postmitotic neurons during human cortical neurogenesis in vitro and in vivo

The function of ASCL1 as a pioneer transcription factor has been established by studying the reprogramming of somatic cells into neurons (Chanda et al., 2014; Raposo et al., 2015; Wapinski et al., 2017; Wapinski et al., 2013), yet the role of endogenous ASCL1 in human cortical development remains unexplored. To investigate this, we modelled human cortical development *in vitro*, by using a two-dimensional (2D) adherent dual SMAD inhibition protocol to promote the differentiation of human induced pluripotent stem cells (iPSCs) into cortical neurons (modified from (Chambers et al., 2009; Shi et al., 2012)). We first characterized the expression of ASCL1 in this model system. We detected expression of *ASCL1* transcripts by qRT-PCR and expression of ASCL1 protein by western blotting at 17 days of differentiation *in vitro* (DIV) (Figure 1A,B), preceding the onset of expression of neuronal markers *MAP2, HUC/D* and *CTIP2* at days 18-20 (Figure S1A). Addition of g-secretase inhibitor DAPT at DIV23 accelerated and synchronized the cell cycle exit and differentiation of neural progenitor cells (Crawford and Roelink, 2007) (Figure S1B,C). ASCL1 RNA and protein levels increased rapidly and peaked 24 hours after DAPT addition (DIV24), before declining and becoming undetectable after DIV30 (Figure 1A,B). Concomitant with the burst in ASCL1 expression at DIV23-24, neural progenitors exposed to DAPT showed a rapid decrease in expression of the cell proliferation marker *MKi67*, upregulation of cell cycle arrest genes *CDKN1C* and *GADD45G*, and strong upregulation of neuronal markers *MAP2, HUC/D* and *CTIP2* (Figure S1A,D).

**Figure 1.**
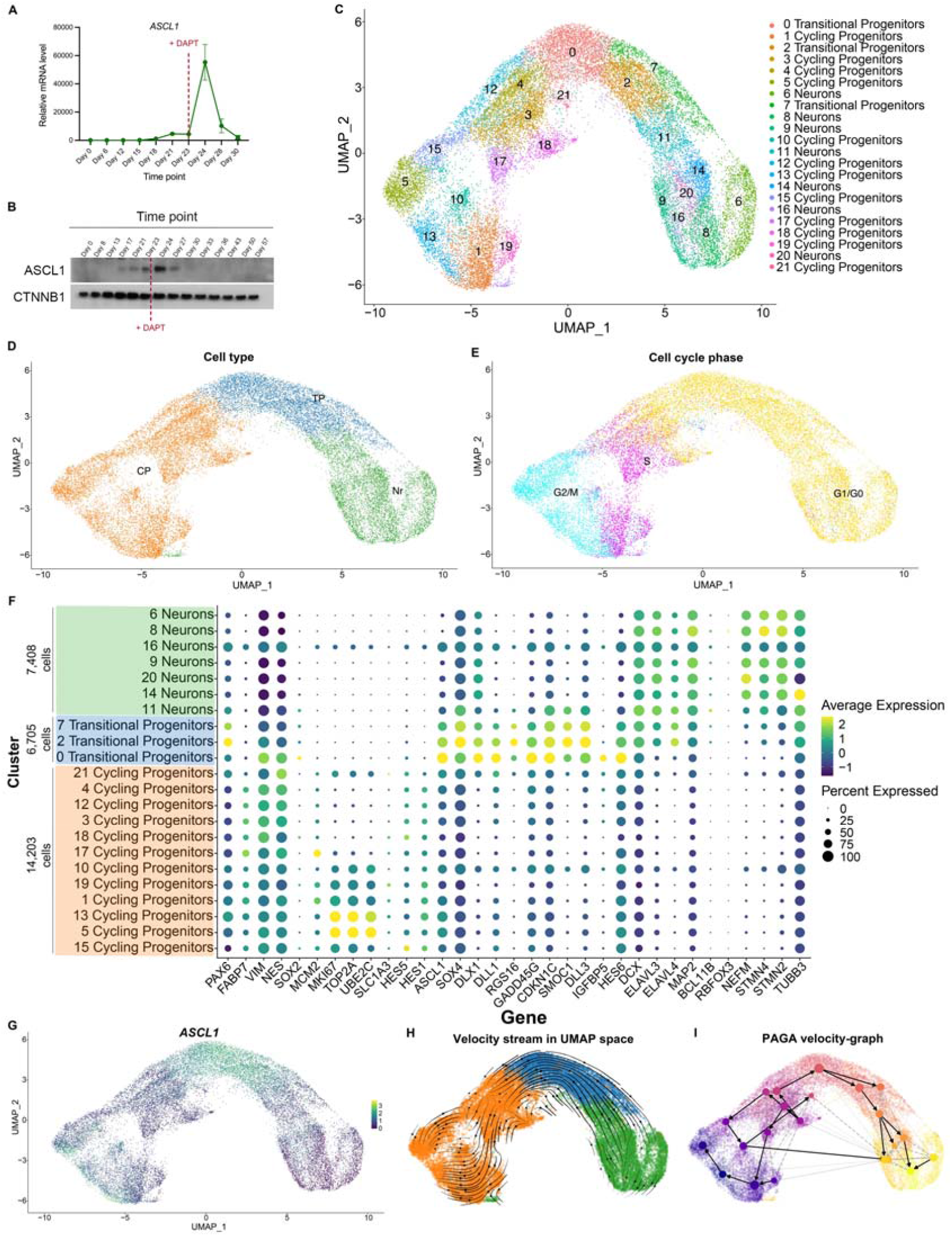
ASCL1 mRNA expression marks a transitional cell population bridging dividing progenitors and postmitotic neurons. **(A)** qRT-PCR analysis of *ASCL1* expression at multiple timepoints during neural differentiation of human iPSCs; mRNA expression relative to DIV0. *ASCL1* shows a transient spike in expression at DIV24, following Notch inhibition (via DAPT addition indicated by a red dashed line) at DIV23. Error bars represent mean ±SEM for three biological replicates. **(B)** Western Blotting shows transient ASCL1 protein increase during neural differentiation. Notch inhibition indicated by a red dashed line. CTNNB1 loading control is included. **(C)** Uniform Manifold Approximation and Projection (UMAP) plot and unsupervised clustering of single-cell transcriptomes from 28,316 cells in DIV24 neural cultures treated with γ-secretase inhibitor DAPT, collected from 3 independent cultures. Dots represent single cells. Colors represent the different clusters identified on the right. **(D)** Clusters from (C) were grouped into 3 broad cellular state clusters based on gene expression of canonical markers (see F). A large cluster of *VIM*+*NES*+ cells, uniquely enriched for *ASCL1* and its transcription targets, lacking in neuronal markers and positioned between a cluster of cycling progenitors (CP) and a cluster of neurons (Nr), was termed ‘Transitional Progenitors’ (TP). **(E)** Predicted cell-cycle phases of cells based on their expression of cell cycle-related genes. Transitional progenitors are found in the G1/G0 phase. **(F)** Dot plot representation of the expression of genes used to annotate cell state identities represented in (D). Dot size indicates proportion of cells in each cluster expressing a gene, shading indicates the relative level of gene expression, according to the key on the right. **(G)** Single gene expression overlayed onto UMAP plot defined in (C) shows *ASCL1* expression is enriched in transitional progenitors. **(H)** RNA velocity vectors projected onto the UMAP plot shows differentiation directionality from Cycling Progenitors to Neurons, through Transitional Progenitors. A second direction was identified that reflected the cell cycle phases. **(I)** A partition-based graph abstraction (PAGA) velocity graph (right), with PAGA connectivities (dashed) and transitions (solid/arrows), shows a similar differentiation trajectory. The size of a node reflects the number of cells belonging to the corresponding cluster.

To further investigate the concomitance of ASCL1 expression with cell cycle exit, we dissected the heterogeneity of neural lineage cell states in iPSC-derived neural cultures by performing a single cell RNA-Seq (scRNA-Seq) analysis in DIV24 cultures treated with g-secretase inhibitor DAPT, using the 10X Genomics Chromium single cell platform. Unsupervised clustering of a dataset of 28,316 cells collected from 3 concurrent differentiated cultures resulted in 22 clusters of cells (Figure 1C, Figure S1E). The clusters were then grouped based on their expression of pre-defined canonical markers of the progenitor state (*SOX2, PAX6, HES5*), proliferation (*MKi67, MCM2*), cell cycle exit (*CDKN1C, GADD45G*) and neuronal differentiation (*HES6, SOX4, DCX, MAP2, ELAVL4, GAD2, SLC17A6)*. This resulted in 3 main cell clusters containing respectively cycling neural progenitors, neurons, and transitional progenitors bridging the two previous clusters (Figure 1D,E,F). The cluster defined as ‘transitional progenitors’ co-expressed markers of progenitors (*VIM, NES*), cell cycle exit (*CDKN1C, GADD45G, HES6*) and early neuronal differentiation (*HES6, SOX4*) with very low levels of proliferation markers (*MKi67* and *MCM2*) and of neuronal markers (*MAP2, NEFM, STNM2, STMN4*; Figure 1F). Thus, this cluster represents progenitors transitioning between mitotic progenitors and differentiating neurons, i.e., likely preceding or just following the last progenitor cell division (Figure 1D,E,F). Differential gene expression analysis confirmed that the transitional progenitor cluster is uniquely enriched for *ASCL1* (Figure 1F,G) and several of its previously reported transcriptional targets including *SMOC1, RGS16*, and *IGFBP5* (Castro et al., 2011; Martynoga et al., 2012), supporting that it is a *bona fide* cell state defined by ASCL1 expression, rather than a mixture of progenitors and neurons. To characterize the cell state transitions in our model of neural differentiation, we predicted directed dynamic transitions between clusters using RNA velocity analysis (with the scVelo tool (Bergen et al., 2020)). The estimated velocity vector streams delineate the direction of cell state transitions with cycling progenitor cells at the apex and a trajectory towards transitional progenitors followed by postmitotic neurons (Figure 1H). Further trajectory inference using PAGA (Partition-based graph abstraction) was employed to estimate connectivity and transition between groups of cells (Wolf et al., 2019) (Figure 1I), which complemented by RNA velocity information supported the directionality of the transitions between cell states. Together, our scRNA-Seq analysis shows that during human neurogenesis, an increase in ASCL1 expression specifically marks a transient population of progenitors differentiating into neurons.

We next sought to further characterize this unique cluster of ASCL1-expressing progenitors identified by scRNAseq, by investigating co-expression of ASCL1 protein with canonical markers of cell identity. Using immunolabelling of DIV24 cultures, we found that ASCL1 was expressed in the granular ‘salt-and-pepper’ pattern characteristic of proneural factors (Kageyama et al., 2008). While all ASCL1-positive cells co-expressed the neural progenitor marker PAX6, none co-expressed the deep layer cortical neuronal marker CTIP2, confirming that ASCL1 is expressed by a subset of progenitor cells and not by differentiated neurons (Figure 2A). Flow cytometry analysis of dissociated DIV24 cultures confirmed this, as shown by the co-expression of ASCL1 with the progenitor marker SOX2 but not with the neuronal marker TUBB3 (Figure 2B). Interestingly, ASCL1 immunolabeling signal was highest in cells bridging progenitors (low TUBB3, high SOX2) and neurons (high TUBB3, low SOX2), suggesting that ASCL1 protein is expressed at its highest levels before the reduction of SOX2 expression and the onset of TUBB3 expression (Figure 2B,D), similar to the *ASCL1* transcripts’ distribution in transitional progenitors in the scRNA-Seq analysis (Figure 1E). Analysis of nuclear DNA content by DAPI staining to infer cell cycle stage, showed a significantly lower proportion of cells in the S-phase of the cell cycle in ASCL1-high compared to ASCL1-low progenitors (Figure 2C,E). This was accompanied by greater proportions of cells in the G0/G1 and G2/M phases in the ASCL1-high population (Figure 2C,E). Taken together, these data suggest that a high proportion of cells from the ASCL1-high population progress through their last division and exit the cell cycle, presumably before differentiating into neurons.

**Figure 2.**
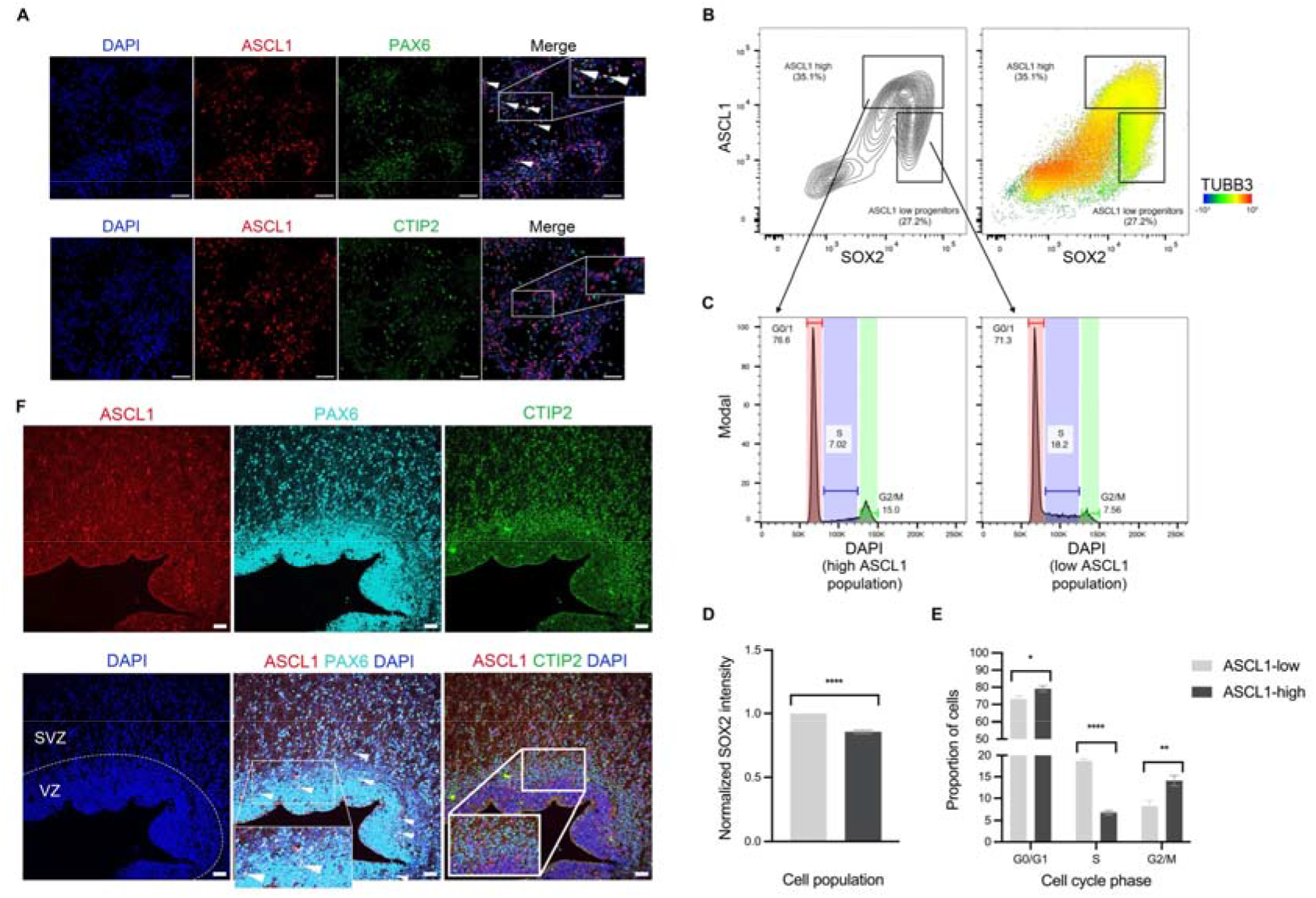
ASCL1 protein expression also marks a transitional progenitor population poised to differentiate. **(A)** Immunofluorescence images of DIV24 neural cultures showing cells co-labelled for ASCL1 and the progenitor marker PAX6 (top, white arrowheads indicate examples), but not for ASCL1 and the neuronal marker CTIP2 (BCL11B, bottom). Bar: 50µm. **(B)** Flow cytometry profiles showing (left) a contour plot of intensities of labeling for ASCL1 and SOX2 in single cells analyzed at DIV24; (right) a dot plot equivalent to the contour plot on the left, pseudocolored according to the level of TUBB3 expression (key on the bottom right). Insets indicate ASCL1-high and ASCL1-low populations. **(C)** DNA content histogram profiles obtained by flow cytometry after DAPI staining for the ASCL1-high (left) and ASCL1-low (right) progenitor populations defined in (B), indicating the cell cycle distribution of cells in these populations. **(D)** Quantification of normalized SOX2 intensities in ASCL1-high and ASCL1-low populations. Error bars represent mean ±SEM for three biological replicates. Unpaired t-test, ****p<0.0001. **(E)** Quantification of the proportion of cells in different cell cycle phases as a percentage of the total cell population (cell cycle analysis by flow cytometry, as analyzed in C), showing significant accumulation of cells in S-phase in the ASCL1-low population and accumulation of cells in G1 and G2/M in the ASCL1-high population. Error bars represent mean ±SEM for three biological replicates. Unpaired t-test, *p<0.05, ** p<0.01, **** p <0.0001. **(F)** PCW 16 fetal brain slice co-immunolabeled for ASCL1, PAX6 and CTIP2, showing co-labelling of ASCL1 with progenitor marker PAX6 (e.g. white arrowheads), but not with deep layer marker CTIP2 (BCL11B). Bar: 50µm. VZ: ventricular zone; SVZ: subventricular zone.

To confirm the relevance of these findings *in vivo*, we investigated ASCL1 expression in the developing human cortex using human fetal brain tissue at post-conceptional week 16 (PCW 16). Immunolabeling of human fetal brain slices for ASCL1 and PAX6 or CTIP2 showed that during human cortical development, ASCL1 expression is also restricted to PAX6-positive progenitors and excluded from postmitotic neurons (Figure 2F), as previously reported (Alzu’bi and Clowry, 2019), and corroborating the findings in our *in vitro* model system of human neurodifferentiation.

### ASCL1 drives the differentiation of cycling progenitors into postmitotic neurons by directly regulating hundreds of genes

Having defined a unique ASCL1-expressing progenitor population consistent with cells exiting the cell cycle, we sought to investigate the role of ASCL1 in that transition. To that end, we generated *ASCL1* knockout (*ASCL1* KO) cells by using CRISPR/Cas9 to create a frameshift-inducing deletion in the same iPSC line (Figure S2A). We genomically screened 3 independent clones for frameshift mutations, differentiated them to DIV24 and performed ASCL1 detection by western blotting. All mutant clones showed an absence of ASCL1 protein even at high chemiluminescent exposure (Figure S2B), thus confirming the generation of *ASCL1*-null mutant cells. To address *ASCL1* function in iPSC-derived differentiating neural cultures, we performed scRNA-Seq on DIV24 cultures of the 3 *ASCL1* KO clones, yielding 35,755 single cells. When we projected this new single cell gene expression dataset onto the reference wild-type UMAP embedding from Figure 1D, we found an increase in the number of cycling progenitors in mutant cells compared with wild-type controls, accompanied by a dramatic reduction in the proportions of transitional progenitors and a near absence of neurons (Figure 3A,B). Moreover, the small number of mutant cells assigned to the transitional progenitors had a different transcriptional signature when compared to control cells (Figure 3C). Because projecting *ASCL1* KO cells onto the predefined WT UMAP embedding forces assignment into the graph based on gene expression pattern, we also independently integrated scRNA-seq datasets for *WT* and *ASCL1* KO transcriptomes and assigned cluster identity based on the previously defined markers (Figure S2C). This analysis corroborated the previous finding of transitional progenitor and neuron depletion in the *ASCL1* KO dataset (Figures S2C,D). When *ASCL1* KO and wild-type cells are integrated, the de novo UMAP representation displays poor clustering of *ASCL1* KO cells with WT cells in the transitional progenitor and neuron clusters, also supporting that the overall transcriptional signatures are different in these two cell types (Figure S2E). Together, these data suggest that loss of *ASCL1* impedes cell cycle exit and differentiation even in the presence of a potent NOTCH inhibitor, resulting in an accumulation of cells in a proliferating and undifferentiated state.

**Figure 3.**
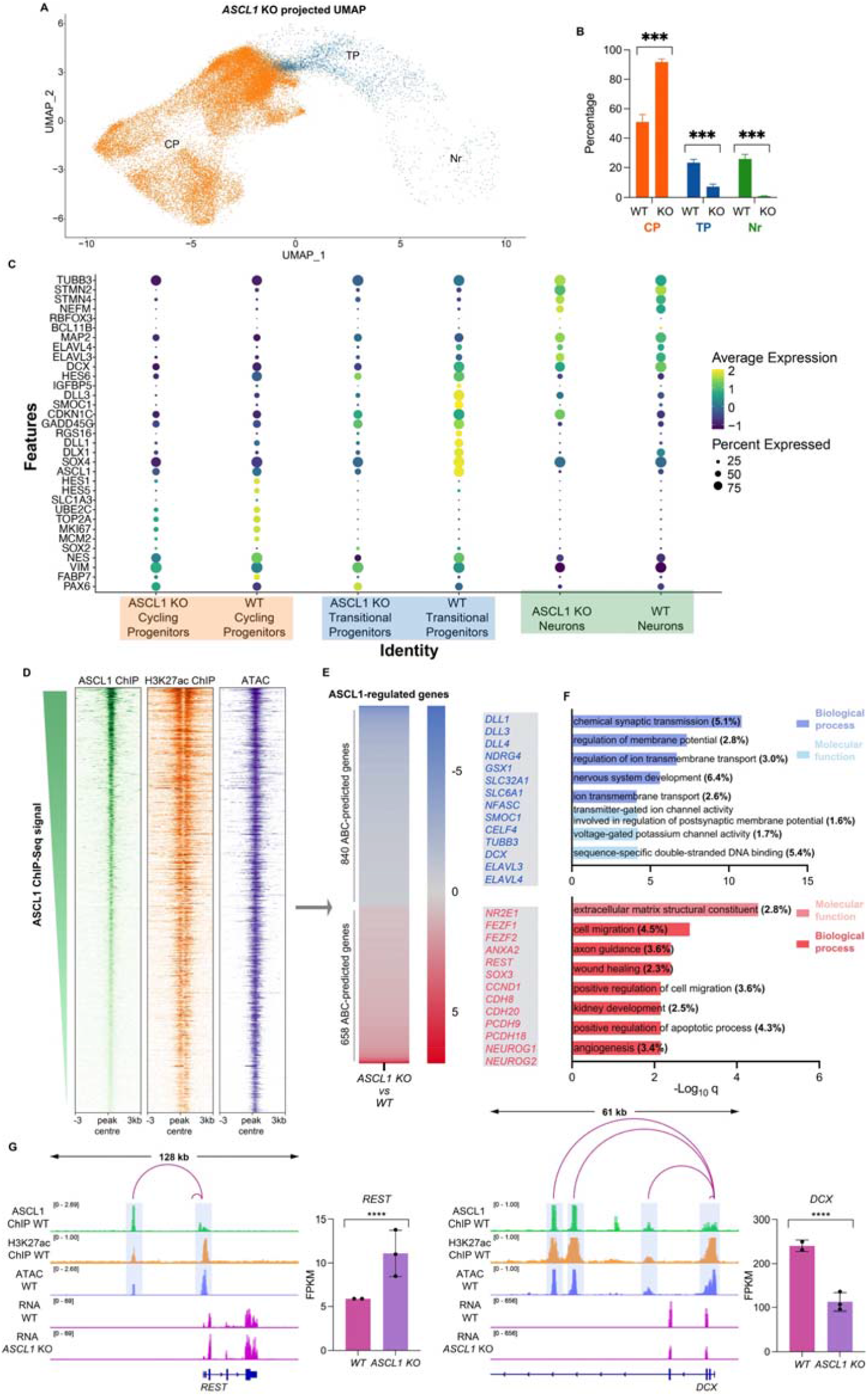
ASCL1 binds to distal regulatory elements of many target genes involved in neural development. **(A)** Single-cell transcriptomes from 35,755 differentiating *ASCL1*-KO neural cultures at DIV24, collected from 3 independent cultures, projected onto the wild-type cells UMAP embedding represented in Figure 1D. **(B)** Relative proportion of major cell clusters from panel (A) in comparison to the cell clusters in Figure 1D. Loss of ASCL1 results in significantly reduced proportions of Transitional Progenitors (TP) and Neurons (Nr) and a significant increase in Cycling Progenitors (CP). Unpaired t-test; ***p<0.001. **(C)** Dot plot representation of the expression of biologically relevant genes in the major clusters from Figure 1D, showing expression differences between WT and *ASCL1* KO cells mapped to those clusters. Dot size indicates proportion of cells in cluster expressing a gene, shading indicates the relative level of expression. **(D)** Heatmaps representing ChIP-Seq coverage for ASCL1 binding sites at genomic regions predicted to be regulatory elements by the ABC algorithm (based on their chromatin accessibility and H3K27ac signature; (Fulco et al., 2019)). **(E)** ASCL1-regulated genes: i.e. genes selected after enhancer-gene regulatory relationship was predicted with the ABC algorithm and where enhancers are identified to be bound by ASCL1 via ChIP-seq, and subsequently the regulated genes were found to be differentially expressed in bulk RNA-Seq analysis of *ASCL1* KO versus wild-type cells (Figure S2). Color coding indicates differential gene expression fold change in *ASCL1* KO vs. wild-type cultures. Illustrative genes for each category are highlighted (listed in Table S1). **(F)** Enrichment of GO biological process terms for the genes downregulated (top, blue) and upregulated (bottom, red) in *ASCL1* KO cells at DIV24, as displayed in panel (E). **(G)** Representative Integrative Genomic Viewer (IGV) tracks of ChIP-Seq, ATAC-seq and bulk RNA-Seq data illustrating examples of ASCL1-occupied active regulatory elements (as predicted by ABC algorithm), targeting genes upregulated (left) or downregulated (right) in *ASCL1* KO versus control cultures at DIV24. Bar plots show mean expression in FPKM for the depicted genes in wild-type and *ASCL1* KO cultures. \*\*\*\**padj*<0.0001.

Because ASCL1 has been shown to drive changes in cell fate through its pioneer transcription factor activity (Chanda et al., 2014; Raposo et al., 2015; Wapinski et al., 2017; Wapinski et al., 2013), we hypothesized the same function could underlie its role in human neurogenesis. Therefore, we investigated *ASCL1* function in transcriptional regulation at DIV24. First, we analyzed transcriptional changes by bulk RNA sequencing of DIV24 cultures in *ASCL1* KO vs control cells. We identified 2562 differentially expressed genes (fold change > 1.5, q-value < 0.05), including 1451 downregulated and 1111 upregulated genes in KO cells (Figure S2F, Supplemental File S1).

We next sought to determine whether these transcriptional changes were due to functional binding activity of ASCL1. First we predicted the genome-wide map of enhancer-promoter interactions specific to our model system (Fulco et al., 2019) using the Activity-By-Contact (ABC) computational algorithm. For this, we characterized the chromatin accessibility landscape by Accessible Chromatin with Sequencing (ATAC-Seq) and the genomic profile of the H3K27ac histone mark (typically present at active enhancers and promoters) by ChIP-Seq in wild-type DIV24 cultures (Figure 3D). Integrating the cell type-specific expression from our wild-type DIV24 RNA-Seq dataset, we were able to predict the enhancer-gene regulatory connections in our human differentiating neural culture system. We identified a repertoire of 23,662 active regulatory elements associated with 10,430 expressed genes (with each gene regulated by one or more elements) (Figure 3G) (Supplemental File S2).

We then characterized the genomic binding profile of ASCL1 by chromatin immunoprecipitation coupled with high-throughput sequencing (ChIP-Seq). Analysis of ASCL1 ChIP-seq at DIV24 revealed 56,100 significant binding sites. The majority of these (70.3%) were intergenic or located in gene introns. The average distance between ASCL1 binding targets and nearest transcription start sites (TSS) was 33.3kb, with most (82.5%) locating more than 1kb from a TSS. Based on the ABC-predicted regulatory landscape, ASCL1 bound 17,049 active enhancer regions in WT DIV24 (Figure 3D). These data are consistent with ASCL1 binding regulating neurogenesis by predominantly binding at distal regulatory elements, in agreement with observations in mouse models of neurodevelopment (Raposo et al., 2015). Having thus defined the regulatory landscape in wild-type cells, we sought to characterize the role of *ASCL1* by investigating the effect of *ASCL1* depletion. From the genes found to be differentially expressed in *ASCL1* KO cells (Figure S2F), ASCL1 binds at least one of the ABC-predicted regulatory elements assigned to 1498 genes (Figure 3E). This repertoire of 1498 herein termed “ASCL1-regulated genes” at DIV24 includes known markers of neuronal differentiation, e.g. neuronal genes *TUBB3* and *DCX* (downregulated in *ASCL1* KO cells) and neuronal transcriptional repressor *REST* (Ballas et al., 2005)(upregulated in *ASCL1* KO cells) (Figure 3E, Supplemental File S3). Gene Ontology analysis of the downregulated *ASCL1* target genes revealed an enrichment of biological processes linked to neuronal differentiation and neuronal activity. Conversely, GO terms associated with upregulated *ASCL1* target genes cells highlighted processes prominent in neural progenitors, including cell adhesion, cell proliferation and inhibition of neuronal differentiation (Figure 3F,G, Supplemental File S3). These experiments therefore establish *ASCL1* as a direct regulator of neural loci during human neurogenesis.

### ASCL1 has pioneer transcription factor activity at a subset of its targets during human neurogenesis

The pioneer transcription factor function of ASCL1 is well established, with evidence for the ability of the overexpressed protein to bind closed chromatin and promote local DNA accessibility, allowing the binding of factors that do not share such a pioneer activity, and regulating cell fate (Chanda et al., 2014; Park et al., 2017; Raposo et al., 2015; Wapinski et al., 2013). However, its pioneer activity in human neurogenesis, and specifically the activity of the endogenous protein, have not been explored. We therefore investigated whether the regulation of the identified *ASCL1*-regulated genes represented canonical features of pioneer transcription factor activity, with a functional interaction between binding of ASCL1 and chromatin accessibility. We first examined how ASCL1 binding correlated with chromatin state in DIV24 cultures by overlapping the wild-type ASCL1 ChIP-Seq and ATAC-Seq datasets. We found that that 31.7% of the 56,100 ASCL1-bound sites in DIV24 cultures were in closed chromatin and 68.3% were in accessible chromatin in the ATAC-Seq dataset, revealing a strong correlation between ASCL1 binding and accessible chromatin at the time point when ASCL1 is highest expressed (Figure 4A). Next, to investigate whether ASCL1 is involved in regulating chromatin structure, we examined chromatin accessibility in DIV24 *ASCL1* KO cultures. We used the ATAC-Seq data to show that among the ASCL1-bound sites in open chromatin in wild-type cells (Figure 3D), 9156 (23.9%) showed changes in accessibility; 46.8% of these showed decreased accessibility in the *ASCL1* KO, and 53.2% regions showed increased accessibility (Figure 4B). Using our pre-determined ABC-predicted regulatory elements and connection with genes they are likely to regulate (Figure 3D), we assigned 460 *ASCL1*-regulated genes to the loci of differential accessibility in *ASCL1* KO vs. wild-type cells. Differential regulatory element accessibility was largely consistent with differential gene expression in the *ASCL1* KO: decreased accessibility was associated with decreased expression in 79.3% ASCL1-regulated genes; conversely, 74.9% of *ASCL1*-regulated genes showed increased accessibility of regulatory elements associated with increased gene expression (Figure 4B). These results are consistent with the notion that pioneer TFs can promote both activation and repression of gene expression depending on secondary factors accessing the opened chromatin (Hosokawa et al., 2018; Zaret, 2020). Our analysis therefore supports that *ASCL1* controls expression of a fraction of its targets by regulating the chromatin accessibility landscape at active regulatory elements, herein termed “*ASCL1*-dependent” genes, thus acting as a pioneer transcription factor.

**Figure 4.**
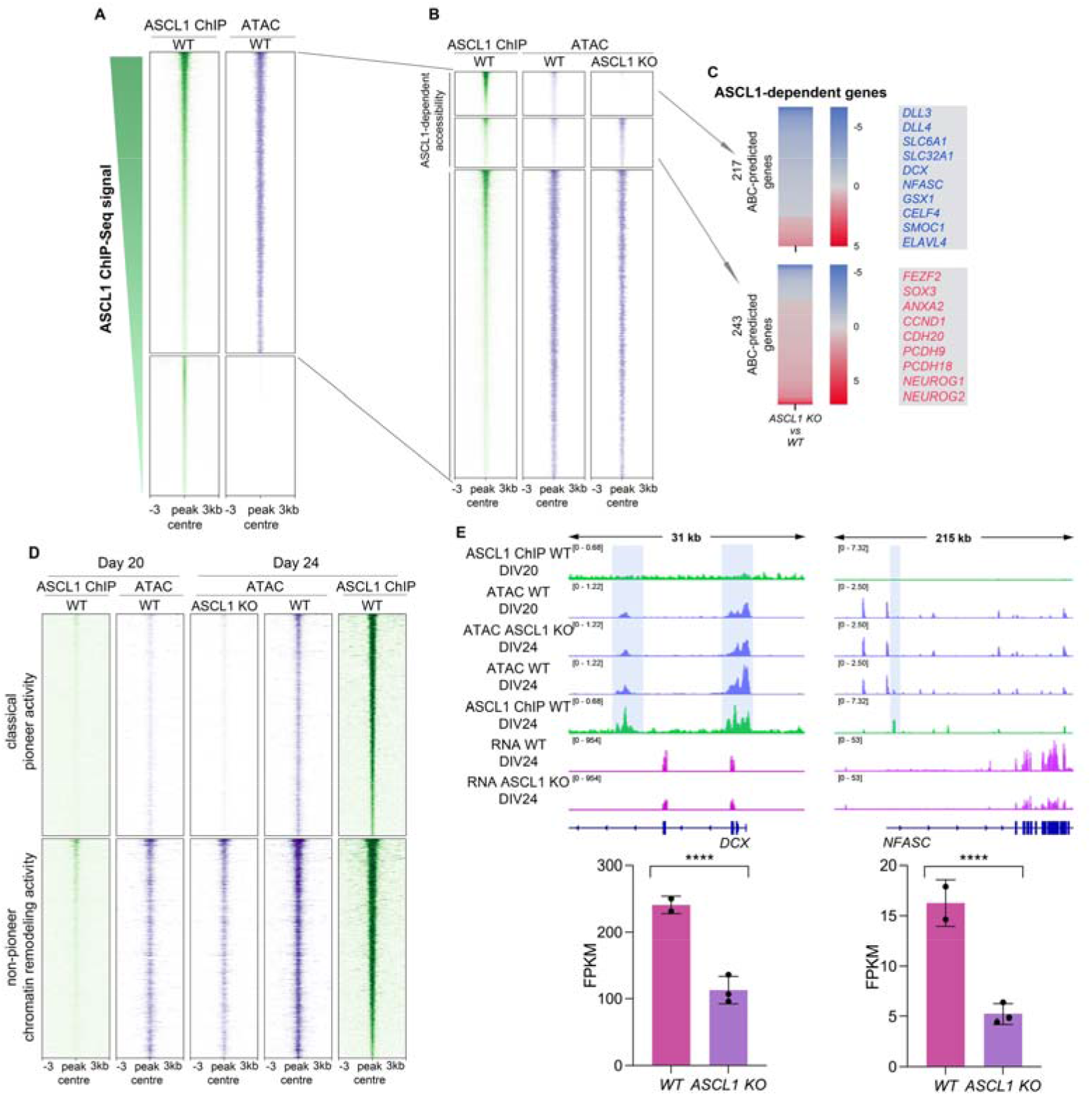
ASCL1 regulates chromatin accessibility through different pioneer and non-pioneer functions. **(A)** Heatmaps showing ASCL1 binding sites identified by ChIP-Seq in open (top, n=38,377) and closed (bottom, n=17,723) chromatin, as determined by ATAC-Seq in DIV24 neural cultures. **(B)** Heatmaps showing open chromatin sites in DIV24 (from A) where ASCL1 regulates accessibility (i.e., “ASCL1-dependent” sites) by opening (top, n=4,288) or closing (middle, n=4896) chromatin, and where ASCL1 binds without regulating accessibility (bottom, n=29,220). **(C)** Heatmaps showing the differential gene expression for ABC-predicted genes associated with regulatory elements at which ASCL1 promotes (top) or represses (bottom) chromatin accessibility (from panel B), and dysregulated in *ASCL1* KO versus wild-type DIV24 cultures. Color coding indicates differential gene expression fold change vs. wild-type culture. Illustrative genes for each category are highlighted. **(D)** Heatmaps showing the changes in chromatin accessibility between wild-type DIV24 cultures (right) and wild-type DIV20 cultures (left) and ASCL1-mutant DIV24 cultures (middle) at ASCL1-dependent sites (from B) where ASCL1 opens chromatin, acting as a classical pioneer transcription factor (top, n=2,155), and where its activity changes accessibility at permissive sites (bottom, n=2,133). **(E)** Representative IGV tracks flanking an ASCL1-dependent gene showing ChIP-Seq and ATAC-Seq profiles at DIV20 and DIV24 (from D) (top). Bottom, bar plots show mean expression in FPKM for the depicted genes in wild-type and *ASCL1*- KO cultures. \*\*\*\**padj* <0.0001.

To gather further evidence for ASCL1 pioneer activity at these ASCL1-dependent sites, we examined the temporal dynamics of chromatin accessibility in differentiating neural cultures and its relationship with ASCL1 binding. We compared chromatin accessibility in wild-type cells at DIV24 with *ASCL1* KO neural cultures at DIV24 (when ASCL1 expression is at its peak), and with wild-type cultures 4 days earlier, at DIV20, when ASCL1 expression is still very low (Figure 1A,B). Because ATAC-Seq is a population-wide genomic assay and likely to reflect both direct and indirect effects of ASCL1 transcriptional activity, we focused our analysis on sites bound by ASCL1 (by ChIP-Seq) in wild-type DIV24 cultures. The first observation was that the accessibility profile of ASCL1 at the pre-bound state at DIV20 mirrored that of the *ASCL1* KO at DIV24 (Figure 4D,E): 74.7% of sites differentially closed at DIV20 vs. wild-type DIV24 were also closed in *ASCL1* KO DIV24 cells, supporting that the chromatin profile of the *ASCL1* KO cultures mirrors the pre-bound state.

We then observed that sites that lost accessibility in *ASCL1* KO cultures fell into two categories: 50.3% of sites exhibited features of interaction with a classical pioneer factor, i.e. open in wild-type cells and completely closed in *ASCL1* KO cells (and similarly in DIV20 progenitors) (Figure 4D, top); the remaining 49.7% sites showed a significant decrease in accessibility (fold change > 1.5, q-value < 0.05) but were not devoid of ATAC signal in *ASCL1* KO cultures, contrary to the first category, i.e., chromatin was already permissive in the absence of ASCL1 (Figure 4D, bottom). This shows that ASCL1 pioneer activity is not required at those sites but its presence increases chromatin accessibility, suggesting that another factor is involved in opening chromatin at those sites and ASCL1 may act as partner to further remodel chromatin (Fernandez Garcia et al., 2019; Henke et al., 2009). To explore this possibility further, we performed motif enrichment analysis separately on the first set of sites (where ASCL1 acts as a classical pioneer) and on the second set (where ASCL1 binds at permissive DNA and is required for further chromatin remodeling). In both categories, the top enriched motif discovered was a bHLH binding motif with the motif core containing the previously reported 5’-CAGCTG-3’ ASCL1 consensus binding sequence (Supplemental File S4) (Raposo et al., 2015). This lack of significant difference in motif analysis suggests that a non-sequence-specific DNA binding factor, e.g., a chromatin remodeling complex may be responsible for opening chromatin at those sites where ASCL1 is found to bind permissive DNA. We identified similar dynamics for the sites where ASCL1 represses chromatin accessibility, though here we found evidence for classical pioneer activity at only 15.6% of sites that are closed vs. open in wild-type DIV24 vs. *ASCL1* KO cultures (Figure S4A, top). This is in keeping with a less prominent role of ASCL1 as a repressive pioneer factor as reported for other pioneer factors (Hosokawa et al., 2018; Wapinski et al., 2017; Zaret, 2020). Overall, 23.5% of all ASCL1-dependent sites are subject to its “classical” pioneer activity in human neural cultures, i.e., binding to closed chromatin and remodeling state, while 68.2% of ASCL1 binding sites are consistent with chromatin remodeling at sites of already accessible DNA, i.e., with a non-pioneer chromatin remodeling activity at those sites (Figure 4D, S4A bottom).

### ASCL1 interacts with mSWI/SNF chromatin remodelers at sites where it acts as a pioneer factor

Given the evidence that ASCL1 acts as a chromatin remodeling factor (with or without pioneer activity) in differentiating human neural cultures, we hypothesized that ASCL1 may regulate chromatin states by interacting with chromatin remodelers, and specifically with ATPase-dependent chromatin remodeling complexes (Zaret and Carroll, 2011). mSWI/SNF or BRG1/BRM associated factor (BAF) complexes are evolutionary conserved ATP-dependent chromatin remodeling complexes with pivotal roles in neural development (Braun et al., 2021; Lessard et al., 2007; Yoo et al., 2009). Because our above data support ASCL1 peak expression coinciding with a progenitor state poised to differentiate, we hypothesized that ASCL1 functionally interacts with mSWI/SNF complexes at this crucial stage of neurogenesis.

In order to test this hypothesis, we first examined the expression patterns of mSWI/SNF subunits and compared them with ASCL1 expression pattern in our *in vitro* model system of neuronal differentiation (Figure 1A,B). Core subunits are expressed at all stages of neural differentiation in these cultures (Figure S3A). Confirming the relevance of the DIV24 timepoint in our experimental model and supporting the previously reported npBAF-nBAF subunit switch in mouse neurogenesis, we observed a decrease in the expression of progenitor-specific subunits ACTL6A, SS18 and DPF1/DPF3, accompanied by an increase in the expression of neuronal-specific subunits ACTL6B, SS18L1 and DPF2/PHF10, as cells differentiated from neural progenitors into postmitotic neurons (Figure S3B). Consistent with our findings of impaired generation of transitional progenitors and neurons in *ASCL1* KO cultures, we also observed in the bulk RNA-Seq dataset a decreased expression of all nBAF-specific subunits in the *ASCL1* KO DIV24 cultures, but no change in the npBAF subunits (Figure S3C). In addition, ASCL1 was expressed in cells containing the npBAF subunit ACTL6A but not the nBAF-specific subunit ACTL6B, consistent with ASCL1 being expressed in neural progenitor cells and not in postmitotic neurons (Figure S3A,D).

Based on the observation that ASCL1-expressing cells (predominantly transitional progenitors) co-express npBAF subunits and our hypothesis above that the two functionally interact, we then investigated if that co-expression reflected a physical interaction between the two. We first performed immunoprecipitation of ASCL1 from a lysate of DIV24 neural cultures followed by western blot analysis (co-IP) and found that ASCL1 co-immunoprecipitates with mSWI/SNF subunits SMARCC1 and ARID1A; reciprocally, immunoprecipitates of SMARCC1 and ARID1A also contained ASCL1 (Figure 5A). We then examined the interaction *in situ* by proximity ligation assay (PLA), conducted in DIV24 control and *ASCL1* KO neural cultures (Figure 5B) and on slices of PCW 16 human fetal cortex (Figure 5C). PLA analysis was conducted with antibodies for ASCL1 and for the mSWI/SNF core subunits SMARCC2 or SMARCB1 as well as the neuronal subunit ACTL6B, and PLA signal was detected for both ASCL1-SMARCC2 and ASCL1-SMARCB1 pairs, but not for ASCL1-ACTL6B negative control pair, both in our *in vitro* model system and *ex vivo* human tissue (Figure 5B,C). Together, these results indicate that ASCL1 physically interacts with mSWI/SNF chromatin remodeling complexes during neural development *in vitro* and *in vivo*.

**Figure 5.**
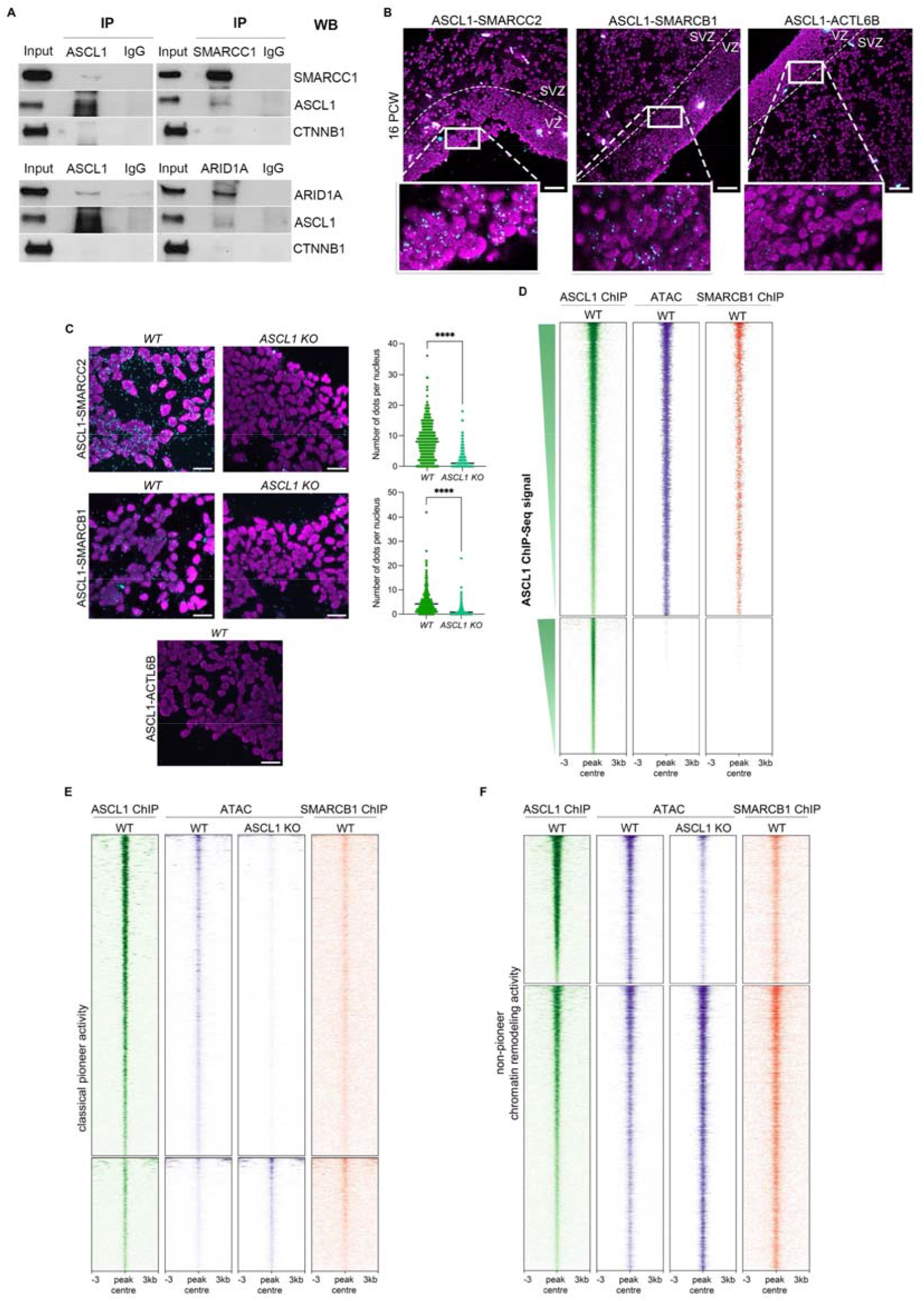
ASCL1 interacts with mSWI/SNF remodelling complexes predominantly at sites where it does not have classical pioneer activity. **(A)** Immunoprecipitation followed by western blot analysis in DIV24 neural cultures showing reciprocal co-immunoprecipitation of ASCL1 and SMARCC1 or ARID1A. CTNNB1 loading control is included. **(B)** Representative immunofluorescence images of Proximity Ligation Assay between ASCL1 and SMARCC2, SMARCB1 or ACTL6B in the human fetal cortex at PCW 16 (B) and in wild-type DIV24 neural cultures (C). Cyan foci indicate PLA amplification signal. Nuclei are shown in magenta (DAPI). **(C)** Number of foci per nucleus were quantified in 3 non-overlapping fields of view. Scale bars, 50µm. No antibody control in Figure S3E. Unpaired t-test; \*\*\*\**p*<0.0001. **(D)** Heatmaps of SMARCB1 binding from ChIP-Seq analysis (right) at all ASCL1 binding sites (left, from Figure 4A, redisplayed for comparison) found in open (top, n=38,377) or closed (bottom, n=17,723) chromatin (middle) in DIV24 neural cultures. Sites co-bound by ASCL1 and SMARCB1 are mostly found in regions of open chromatin. **(E**,**F)** Heatmap profiles of SMARCB1 binding (right) in DIV24 neural cultures at ASCL1-dependent sites (left, from Figure 4, redisplayed for comparison) where (E) ASCL1 acts as a classical pioneer factor binding closed chromatin (ATAC signal below threshold in *ASCL1* KO cultures, middle right) and (F) sites where ASCL1 binds permissive chromatin (non-pioneer chromatin remodeler; ATAC signal shows open signal in *ASCL1* KO cultures but ASCL1 cause changes in accessibility, middle right vs. middle left). SMARCB1 is enriched at sites where ASCL1 binds open chromatin and changes accessibility (F, top and bottom).

To investigate if this physical interaction between ASCL1 and mSWI/SNF complexes is relevant for co-regulation of epigenetic states, we first determined the genome-wide binding profile of the mSWI/SNF core subunit SMARCB1 in DIV24 neural cultures by ChIP-Seq. We found that 90.3% SMARCB1 binding sites overlapped with nucleosome-depleted regions previously identified by ATAC-seq (Figure S4C) consistent with its role in chromatin remodeling (Bao et al., 2015; Iurlaro et al., 2021; Schick et al., 2019). Assuming that the physical interaction between ASCL1 and mSWI/SNF is involved in chromatin regulation, we then overlapped ASCL1 (Figure 4A) and SMARCB1 binding sites: 54.9% of the ASCL1 targets in open configuration at DIV24 also displayed SMARCB1 binding, while only 5.8% ASCL1 targets in closed configuration in DIV24 were co-bound by SMARCB1, indicating that co-recruitment relates to accessible chromatin (Figure 5D). In addition, motif analysis revealed that the motif enriched at the largest number of targets (52.2%) includes the 5’-CAGCTG-3’ ASCL1 consensus binding sequence at its core (Raposo et al., 2015)(Supplemental File S4), suggesting the sequence binding specificity of the cooperative pair is conferred by ASCL1. Conversely, the ASCL1 motif is not identified in sites bound only by SMARCB1.

Because we had identified two classes of ASCL1-dependent loci, where ASCL1 classical pioneer activity is required, and where ASCL1 acts at a non-pioneer remodeling factor, changing chromatin state on already permissive DNA, respectively (Figure 4D), we hypothesized that the requirement of ATP-dependent co-factors might differ between the two classes. Indeed, at sites where ASCL1 acts as a classical pioneer factor, we identified SMARCB1 ChIP-Seq co-binding at only 12.7% of them (Figure 5E). Conversely, at sites where ASCL1 affects accessibility without the classical features of a pioneer factor, 53.1% of them are co-bound by SMARCB1 (Figure 5F). Moreover, examining the SMARCB1 ChIP-Seq data, we found that at most of the 46.9% remaining sites, there is some SMARCB1 ChIP-Seq signal that is below threshold for peak calling. With these data, we conclude that ASCL1 and mSWI/SNF interact at sites where both engage with DNA at open chromatin sites, and that this interaction is more predominant at sites where ASCL1 lacks classical pioneer activity.

### ASCL1 works in concert with the mSWI/SNF ATPase complexes to remodel chromatin and regulate gene expression during human neurogenesis

Given that mSWI/SNF is enriched at ASCL1-dependent sites, particularly at sites where ASCL1 is not required to open chromatin but acts to increase accessibility, we hypothesized that ASCL1 requires mSWI/SNF complexes to regulate chromatin accessibility at those sites. To address this, we investigated the effect of suppressing mSWI/SNF activity on *ASCL1* function. We first chose to acutely eliminate the mSWI/SNF complexes by simultaneous CRISPR/Cas9-mediated knockout of the *SMARCC1* and *SMARCC2* core subunits in DIV21 cultures (preceding the ASCL1 expression peak) with analysis of the mutant cultures 3 days later at DIV24. Elimination of both core subunits (Figure S4D) resulted in the depletion of other mSWI/SNF subunits, likely reflecting destabilization of the entire mSWI/SNF complex in the absence of its core subunits (Figure S4E), in agreement with reports in other models (Mashtalir et al., 2018; Schick et al., 2019). However, in addition to poor cell viability with almost complete depletion of mSWI/SNF complexes, ASCL1 expression itself was downregulated in SMARCC1/C2-mutant cells (Figure S3F), limiting the value of this model for investigating the role of mSWI/SNF-ASCL1 interaction.

We thus chose the alternative approach of using the small molecule BRM014 to inhibit the catalytic activity of the two ATPases of mSWI/SNF complexes, SMARCA2 and SMARCA4 (Iurlaro et al., 2021; Papillon et al., 2018). We exposed DIV22 neural cultures to 10mM BRM014 and harvested the cultures for analysis 48hrs later, at DIV24 (one day after NOTCH inhibition, at ASCL1 expression peak). ASCL1 expression was not affected by exposure to BRM014 (Figure S4G). We first investigated how BRM014 treatment affected mSWI/SNF complex enzymatic activity by examining its effect on chromatin accessibility using ATAC-Seq in BRM014-treated and control cells. We found that BRM014-treated cells presented reduced chromatin accessibility at 3,664 genomic sites and increased accessibility at 277 sites (q < 0.5), with 68.6% of the changes in accessibility occurring at SMARCB1 ChIP-Seq binding sites. These results were in keeping with the anticipated effects of inhibiting mSWI/SNF enzymatic activity without compromising ASCL1 expression, indicating this is a suitable system to investigate mSWI/SNF-ASCL1 interactions.

To address if ASCL1 requires mSWI/SNF to regulate chromatin states, we focused our attention on ASCL1-dependent sites where ASCL1 binding significantly affects chromatin accessibility (see Figure 4). In 41.6% of the ASCL1-dependent sites in BRM014-treated cells, we identified a concordant change in chromatin state: 54.2% are sites that show decreased accessibility in both *ASCL1 KO* (Figure 4B, top; 4D) and BRM014-treated cells; conversely, 30.6% are sites with increased accessibility in both *ASCL1 KO* (Figure 4B, middle, S4A) and BRM014-treated cells. For these ASCL1-dependent sites where chromatin accessibility changes are concordant in both conditions, 66.3% are also bound by SMARCB1 on ChIP-Seq in wild type cells (Figure S4H). Moreover, when we plotted the accessibility changes at ASCL1-dependent sites in *ASCL1* KO cells and BRM014-treated cells, we found a positive correlation (r=0.395) between the two conditions (Figure S4H). We found a similar correlation when comparing the transcriptional effects of both conditions (Figure S4I) indicating that the interaction of ASCL1 with mSWI/SNF is relevant to downstream transcriptional activity.

Hence, we reasoned that the ASCL1-mSWI/SNF interaction inducing changes to chromatin structure in transitional progenitors would have relevance to the regulation of neural loci. We used once again the active enhancer-promoter regulatory relationships predicted by the ABC algorithm (Fulco et al., 2019) to assign the relevant genes co-regulated by ASCL1 and mSWI/SNF ATPase activity (co-bound and with differential accessibility in both *ASCL1* KO and BRM014-treated cells). We identified 1259 co-bound predicted regulatory elements that control the expression of 141 genes during human neurogenesis, corresponding to 30.7% of the *ASCL1*-dependent genes and including important genes involved in neural development, e.g. *DLL3, MYT1, STMN1, SHANK1* activated by both partners (Figure 6C). Specifically, for the 3315 ASCL1-dependent sites where ASCL1 binds permissive DNA (Figure 4D), 53.1% are co-regulated by mSWI/SNF activity (corresponding to 1195 predicted regulatory elements assigned to 132 genes), whereas for the 2915 ASCL1 dependent sites where ASCL1 acts as a classical pioneer TF, we find co-dependency for only 12.7% (64 predicted regulatory elements assigned to 9 genes). These results indicate that cooperative binding of ASCL1 and ATPase active mSWI/SNF is predominantly at sites where chromatin is already partially or transiently opened.

**Figure 6.**
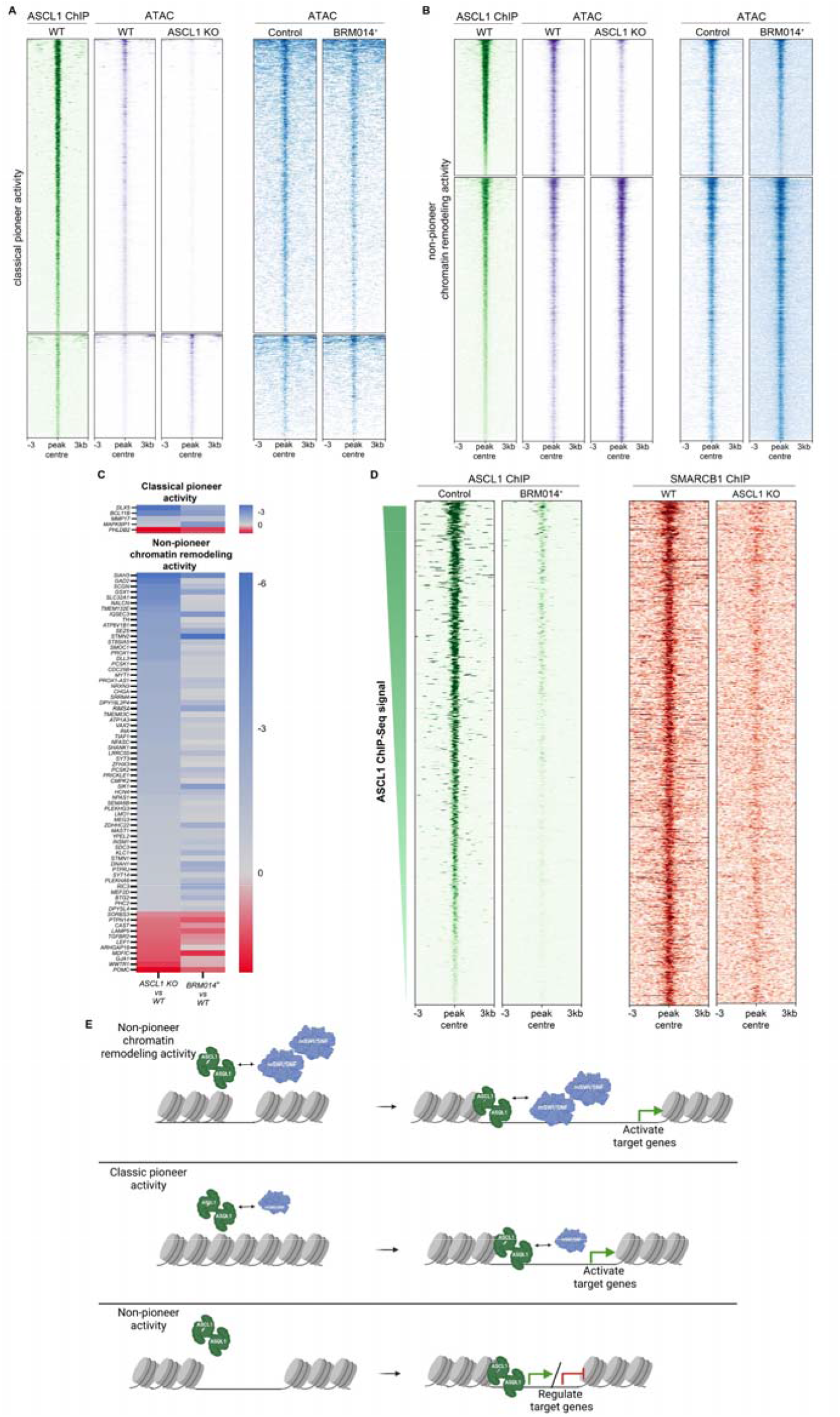
ASCL1 works in concert with mSWI/SNF complexes to remodel chromatin and regulate gene expression. **(A, B)** Heatmaps showing the effect of BRM014-treatment on chromatin accessibility at ASCL1-bound sites where ASCL1 displays classical pioneer activity (A) or non-pioneer chromatin remodeling activity (B). The ATPase activity of mSWI/SNF complexes is mainly required at sites of non-pioneer activity (B). ASCL1 ChIP-Seq (Figure 5E) is included for reference. **(C)** Top: Heatmap of differential gene expression for genes dysregulated in both *ASCL1* KO and BRM014-treated cultures, and whose ABC-predicted regulatory element(s) are associated with ASCL1-SMARCB1 co-bound sites where both are required to increase accessibility, and where ASCL1 acts as a classical pioneer factor. Bottom: heatmap of differential gene expression for genes dysregulated in both *ASCL1* KO and BRM014-treated cultures, and whose ABC-predicted regulatory element(s) are associated with ASCL1-SMARCB1 co-bound sites where both are required to increase accessibility, and ASCL1 acts as a non-pioneer chromatin remodeler. **(D)** Heatmaps profiling ASCL1 and SMARCB1 binding at ASCL1-mSWI/SNF dependent sites where the interaction is associated with open chromatin, showing that interfering with mSWI/SNF ATPAse activity (BRM014 treatment) reduces ASCL1 binding and reciprocally, eliminating ASCL1 (*ASCL1* KO) reduces SMARCB1 binding. **(E)** Diagram illustrating the proposed mechanism for the ASCL1-mSWI/SNF recruitment dynamic: the two interacting partners depend on each other for binding at regulatory elements where they regulate accessibility. Created with BioRender.com.

Having identified co-binding by ASCL1 and mSWI/SNF and co-regulation of chromatin accessibility, we then asked whether chromatin remodeling activity of one factor at the co-bound sites is required for binding of the other, or whether each factor can bind independently of the other. To investigate this, we performed reciprocal SMARCB1 and ASCL1 ChIP in *ASCL1* KO and BRM014-treated cells, respectively. At 92.7% of these putatively co-regulated sites, we found evidence for a significant decrease or complete absence of DNA binding of the partner (Figure 6C). Residual binding of SMARCB1 is consistent with the limited chromatin accessibility observed in *ASCL1* KO cells at sites of non-pioneer remodeling activity (Figure 4B). We thus propose a mechanism of cooperative binding of ASCL1 and mSWI/SNF at active distal regulatory elements resulting in co-regulation of chromatin remodeling and gene regulation (Figure 6C,D).

## Discussion

We used genome-wide epigenetic analyses to investigate the role of ASCL1 in human neurogenesis. De-repression of ASCL1 induced by NOTCH inhibition induces terminal differentiation (Park et al., 2017) by direct binding of ASCL1 to DNA, activating the neurogenic program. The ASCL1-dependent progenitors we identified are a transient population of cells that rely on ASCL1 for cell cycle exit and neurodifferentiation.

Although a progenitor population has already been termed “transitional progenitors” in an earlier study (Earley et al., 2021), the neuronal differentiation protocol was very different and the cell cluster, enriched in neurogenic transcription factors including but not defined by ASCL1, likely represents a very different cell stage. ASCL1 regulates hundreds of genes in neural differentiation cultures, and we find evidence for direct binding of distal regulatory elements in transitional progenitors. Investigating ASCL1’s pioneer transcription factor function, we identified two groups of regulated loci: those where ASCL1 acts as a classical pioneer TF, and another where it binds permissive DNA and is required to promote further chromatin changes. This latter group is reminiscent of the “non-classical pioneer factor” class proposed by Minderjahn et al. (Minderjahn et al., 2020) to define PU.1, which is unable to access nucleosomal target sites, but when overexpressed can remodel chromatin and redistribute partner TFs in a mSWI/SNF complex-dependent manner. Here, we use the term “non-pioneer chromatin remodeling activity” to refer to the role of ASCL1 at sites where it affects chromatin state after binding already open DNA. The finding that mSWI/SNF function is also involved in this activity suggests similar mechanisms may be at play in the two systems.

Cooperative binding of chromatin remodeling complexes and pioneer transcription factor has come under particular attention in recent years. mBAF SWI/SNF has been implicated as an interactor with other pioneer factors, such as OCT4, GATA1, GATA3 and KLF4 (Hu et al., 2011; King and Klose, 2017; Moonen et al., 2022; Takaku et al., 2016). In mouse ES cells, SMARCA4 ATPase subunit of mSWI/SNF is required at OCT4-dependent sites to regulate chromatin accessibility at distal regulatory elements and to allow binding of other pluripotency-associated transcription factors (King and Klose, 2017). Pioneer factor KLF4 and SMARCA4 containing SWI/SNF complexes co-occupy active enhancer regions in endothelial cells, where KLF4 regulates chromatin accessibility of vasculo-protective genes in response to laminar sheer stress (Moonen et al., 2022).

We identify a cooperative function of bHLH transcription factor ASCL1 and mSWI/SNF chromatin remodeling complexes at distal regulatory elements in regulating chromatin structure and in promoting neurogenesis at a key transitional stage (Figure 5). We find evidence for co-dependent DNA binding, with interference with one binding partner affecting binding of the other at co-regulated sites (Figure 6). Cooperative interaction between ASCL1 and ATPase-active SWI/SNF is greatest at regulatory elements where ASCL1 acts as a non-pioneer chromatin remodeler (53.1% of sites). Conversely, at loci where ASCL1 acts as a classical pioneer factor, although a subset of the loci is co-bound by mSWI/SNF (12.7%), they do not require its presence to elicit changes in chromatin structure. We observe that interfering with chromatin remodeling (*ASCL1* KO or mSWI/SNF activity inhibition) results in significant reduction in binding of the cooperative partner, though residual binding of mSWI/SNF in the absence of ASCL1 is presumably sufficient for some degree of accessibility. This suggests that the ASCL1-mSWI/SNF interaction may act to stabilize mSWI/SNF at its targets, and that in the absence of ASCL1 mSWI/SNF releases from DNA more readily.

Using molecularly defined active regulatory regions (i.e with ATAC-Seq and H3K27ac ChIP-Seq signals), we computationally defined enhancer-gene pairs regulated by the ASCL1-mSWI/SNF interaction. Genes co-regulated by this interaction and where ASCL1 acts as a non-pioneer chromatin remodeler were enriched for ontology terms associated with cell differentiation, brain development and neurotransmitter processes, supporting a role in directing neural differentiation. There are fewer co-regulated genes where ASCL1 acts as a classical pioneer TF, yet these include important neuronal genes, including *BCL11B* (Supplemental file 3), encoding transcription factor BCL11B/CTIP2 expressed in deep layer cortical neurons (Simon et al., 2020). It is likely that similar to OCT4-SMARCA4 interaction regulating pluripotency (King and Klose, 2017), the pioneer ASCL1-SWI/SNF interaction may facilitate binding of other TFs important for neural differentiation, though further studies would be required to confirm that.

One possibility for the distinct requirements of SWI/SNF as an interacting partner at classical pioneer and non-pioneer sites of chromatin remodeling is that different loci may have architectural constraints that either require or limit ASCL1-DNA interaction. ASCL1 in an heterodimer with bHLH E-proteins, binds an E-box motif with degenerate central two nucleotides (5’-CANNTG-3’) (Fernandez Garcia et al., 2019; Soufi et al., 2015). We did not find evidence for differences in binding motifs at sites of classical and non-pioneer chromatin remodeler activity, therefore we speculate that as per the previously reported ASCL1 pioneer function (Wapinski et al., 2013), cooperative binding between ASCL1 and mSWI/SNF may be determined by factors such as histone modifications, which we do not explore here. A limitation of the study is that molecular characterization of the system is not performed at the single cell level apart from scRNA-Seq. Since ASCL1 is expressed primarily in transitional progenitors, we can assume the ChIP-Seq binding profiles are specific to these cells. However the ATAC-Seq profile reflects the whole cultured cell population, and moreover, as with other DNA accessibility techniques, ATAC-Seq does not perform well on detection of partially unwrapped nucleosomes (Sung et al., 2014). Thus it is possible that the non-pioneer chromatin remodeler ASCL1 repertoire and cooperative binding of mSWI/SNF at those sites represent heterogeneous states that will be more readily deciphered with higher resolution analysis (Buenrostro et al., 2015; Hainer et al., 2019; Harada et al., 2021). As we find cooperative binding for only a small subset of ASCL1 classical pioneer targets, we note that we did not investigate other chromatin remodelers than mSWI/SNF which may act with ASCL1 in regulating chromatin structure at sites of classical pioneer activity.

The use of a constitutive knock-out of ASCL1 rather than conditional ablation may also be a limitation of the study. However, the observation that the chromatin accessibility landscape of knockout cells at ASCL1 peak expression (DIV24) is similar to that of wild-type cells at a timepoint preceding significant ASCL1 expression (DIV20; pre-bound state) suggests that constitutive loss does not have a significant effect prior to NOTCH inhibition and the sharp increase in ASCL1 expression at DIV23-24. Conversely, while we ablated mSWI/SNF enzymatic activity preserving protein expression (Iurlaro et al., 2021; Papillon et al., 2018), we cannot exclude ATPase-independent roles for mSWI/SNF, which might be required at sites of ASCL1 classical pioneer activity bound by mSWI/SNF (Jordán-Pla et al., 2018). Having focused our attention on the cooperative functions of ASCL1 and mSWI/SNF, further investigations are also required to explore whether the roles of mSWI/SNF in the transition from neural progenitors to neurons in mouse neurogenesis are replicated in humans (Staahl et al., 2013; Yoo et al., 2009), and whether these roles reflect predominantly mSWI/SNF’s interaction with ASCL1, or also interactions with other neurogenic factors.

Our findings of a regulatory interaction between ASCL1 and mSWI/SNF chromatin remodelers in controlling the neurogenic process may have implications beyond ASCL1’s role in neural development. Glioblastomas respond to overexpression of ASCL1 by terminally differentiating proliferating cells, thus restricting tumor expansion (Park et al., 2017). The identification of mBAF SWI/SNF dependency at neurogenic regulatory loci may indicate a targetable vulnerability in these cancers.

## Methods

### iPSC Cell culture and differentiation

Experiments were performed using the Human Induced Pluripotent Stem Cell Initiative (HipSci, www.hipsci.org) Kolf2C1 line (clone C1 of parental line HPSI0114ikolf2), kindly gifted by William T. Skarnes at the Wellcome Sanger Institute. The stem cells were maintained at 37°C, 5% CO2 under feeder-free conditions on Geltrex (ThermoFisher Scientific) coated plasticware (Corning) in E8 media (ThermoFisher Scientific) + 100U/ml Penicillin/Streptomycin (ThermoFisher Scientific).

The iPSCs were differentiated into cortical neurons in two-dimensional adherent cultures using a dual SMAD inhibition protocol (modified from (Chambers et al., 2009; Shi et al., 2012)). Briefly, the iPSCs were washed with dulbecco’s phosphate buffered saline (DPBS, ThermoFisher Scientific), dissociated in 0.5mM ethylenediaminetetraacetic acid (EDTA) pH 8.0 (Invitrogen) in DPBS and plated on Geltrex-coated plates in E8 in a 2:1 ratio on day -1. After 24 hours (day 0), the culture reached 80-100% confluency, at which point the medium was replaced with neural induction medium (N2B27, 1:1 mixture of N2 medium and B27 medium (composition listed in Supplemental Table 1) supplemented with 10mM SB31542 (Abcam) and 10nM LDN193189 (StemCell Technologies). The neural induction medium was replaced daily for 7 days. On day 7, the neuroepithelial cells were washed with Hanks’ Balanced Salt Solution (HBSS, ThermoFisher Scientific), detached using Accutase (Sigma-Aldrich) and replated on Geltrex-coated plates in N2B27 with 10mM Y-27632 ROCK inhibitor (Tocris). The ROCK inhibitor was removed after 24 hours (day 8) and the neural induction medium replaced daily until day 12. On day 12, the neural progenitors were dissociated in Accutase and replated on Geltrex-coated plates in N2B27 with 10mM Y-27632 ROCK inhibitor. ROCK inhibitor was removed on day 13 and cells were maintained in daily-changed N2B27 until day 23 (with 1:2 passaging on days 15-16 and 19-20 as already described for days 7 and 12). On day 23, the medium was changed to B27 only supplemented with 10mM DAPT (Cambridge Bioscience). On days 25 and 27, half of the media was replaced with fresh B27 supplemented with 10mM DAPT. From day 30 onwards, the neurons were maintained in B27 medium only, replacing only half of the B27 media volume every week.

Because the Kolf2C1 line was subsequently identified as harboring a somatic ARID2 frameshift mutation (Skarnes et al., 2019), we performed the neural differentiation protocol above on Kolf2C1, the gene-corrected derivative KOLF2.1J (kindly gifted by William T. Skarnes, The Jackson Laboratory for Genomic Medicine, as well as independent HipSci line Kucg2 (HPSI0214i-kucg_2), and analyzed genome wide transcriptomes by RNAseq at DIV24 where most of our experimental work is conducted. We found that the corrected KOLF2.1J and Kolf2C1 have similar transcriptomes, clustering tightly together and apart from the Kucg2 line on principal component analysis (Figure S5). This finding indicates that transcriptional differences attributable to the single gene change were minimal in comparison to those induced by genetic background (Kilpinen et al., 2017) and we thus proceeded with the Kolf2C1 line for which our neuralization protocol had been optimized.

### Generation of ASCL1 knockouts

ASCL1 knockout Kolf2C1 cells were generated with CRISPR/Cas9 methods modified from ref. (Bruntraeger et al., 2019), and using two previously validated *ASCL1* crRNA sequences (Liu et al., 2016). Briefly, wildtype Kolf2C1 cells were supplemented with 10mM ROCK inhibitor 1 hour prior to nucleofection. For each *ASCL1* crRNA, 200mM crRNA and 200mM tracrRNA (IDT) were mixed in IDT duplex buffer (IDT), in a 1:1:2.5 ratio then hybridized by heating to 95°C for 2min before returning to room temperature (r.t.) to form sgRNAs. The 2 sgRNAs were then combined 1:1, and 5ml of this mixed with 20ug Alt-R® S.p. HiFi Cas9 Nuclease V3 (IDT) and incubated at r.t. for 30mins to form Cas9 ribonucleoprotein (RNP) complexes. Concurrently, cells were collected with accutase supplemented with 10mM ROCK inhibitor and washed twice in DMEM F-12 (ThermoFisher Scientific) then strained through a 40mm cell-strainer. 10^6^ cells were then resuspended in 100ml P3 buffer (Lonza) and mixed with the prepared Cas9-RNPs and 5ml EP enhancer (IDT). This was then nucleofected on an Amaxa 4D Nucleofector using program CA137 (Lonza) and plated into a Synthemax-coated (Sigma-Aldrich) 6-well in E8 + ROCK inhibitor. After recovery, cells were plated at single-cell density to obtain colonies of single clones using CloneR (Stemcell Technologies)-supplemented media, following manufacturer’s instructions, and manually picked into 96 well plates. Single clones were screened for desired edits by targeted amplicon high-throughput sequencing (MiSeq) using previously published PCR primers (Liu et al., 2016) with appended Illumina sequencing adaptors and barcodes. Clones with biallelic deletions inducing a frameshift (134 bp deletion, ENST00000266744.4:c.92_225del, ENSP00000266744.3:p.Phe31TrpfsTer81) were expanded and knockout was confirmed by western blotting.

### RNA extraction, cDNA synthesis, and qRT-PCR

Samples were collected in RLT lysis buffer (Qiagen) added directly to the DBPS-rinsed cell culture plates (ThermoFisher Scientific). RNA was then extracted using the RNeasy Micro Kit (Qiagen) according to the manufacturer’s protocol, with 15 minute on-column digestion with RNase-free DNase I (Qiagen) to eliminate genomic DNA. RNA was converted to cDNA using the Maxima First Strand cDNA Synthesis kit with dsDNase (ThermoFisher Scientific), using reverse transcriptase with a mix of random hexamer and oligo(dT) 18 primers, according to the manufacturer’s protocol.

qRT-PCR reactions were prepared with Taqman Universal qRT-PCR Master Mix (ThermoFisher Scientific) and commercially designed primer probes (Supplemental Table 2) following the manufacturer’s protocol. Reactions were prepared in triplicate and run on the Lightcycler 480 II thermal cycler (Roche, Switzerland). Data were exported from proprietary Roche software and subjected to statistical testing in Excel (Microsoft, USA) based on the -2^−DDCt^ or Livak method (Livak and Schmittgen, 2001). Briefly, gene products from the three technical replicates were averaged, followed by normalization against *HPRT* or *UBC* to generate the DCt values, which were then compared to the day 0 values to generate the DDCt values. qRT-PCR analysis was performed on samples obtained from a minimum of three independent experiments. Data were graphed using GraphPad Prism software.

### Western blotting

Cells were washed with DPBS and lysed in Pierce IP lysis buffer (ThermoFisher Scientific) supplemented with 1X Protease inhibitor cocktail (ThermoFisher Scientific), 1X EDTA (ThermoFisher Scientific) and 1X Phosphatase inhibitor cocktail (ThermoFisher Scientific). Cells were scraped off the plates and lysed at 4° for 20 minutes under rotation, followed by centrifugation at 17,000 x g for 20 minutes. Supernatant was collected in a new tube and stored on ice for quantification. A bicinchoninic acid assay (Pierce BCA Protein Assay Kit, ThermoFisher Scientific) was used to quantify the protein extract supernatant according to manufacturer’s protocol. Bovine serum albumin (BSA, ThermoFisher Scientific) was used to generate a standard curve and color change was then quantified using the EnSight multimode plate reader (Perkin Elmer, USA) and analyzed using the proprietary software. Following quantification, protein was stored at -80°C until analyzed by western blot or subjected to immunoprecipitation. Samples were then prepared for sodium dodecyl sulphate polyacrylamide gel electrophoresis (SDS-PAGE) by dilution with 2X (Sigma) or 5X (in house made) Laemmli sample buffer and incubation at 95°C for 5 minutes. Denatured samples were run on 4-15% polyacrylamide gels (Bio-Rad) in 1X Tris-glycine-SDS (TGS) running buffer (Bio-Rad) at 120-130V. A polyvinylidene fluoride (PVDF) membrane (Bio-Rad) was used for sample transfer using the Trans-Blot Turbo Transfer System (Bio-Rad) before blocking in 5% milk (Marvel) in Tris Buffered saline Tween (TBS-T, Bio-Rad) for 60 minutes. Membranes were incubated with primary antibodies (Supplemental Table 3) diluted in 5% milk (Marvel) in TBS-T overnight at 4°C with rocking. The next day, membranes were washed in TBS-T, followed by incubation in secondary antibodies (Supplemental Table 3) diluted in 5% milk in TBS-T at r.t. for 60 minutes. TBS-T washes were performed again and signal was generated using enhanced chemiluminescence (ECL) substrate (Amersham) as per manufacturer’s instructions. A Hyperfilm ECL (Amersham) was developed for signal detection in a dark room.

### Co-immunoprecipitation

For co-immunoprecipitation (co-IP) experiments, primary antibodies (Supplemental Table 3) were added to the protein supernatants and incubated with rotation for 2 hours at 4°C. At the same time, Sepharose coupled with protein G (Sigma) was blocked with 5% BSA (Sigma) in precooled DPBS (ThermoFisher Scientific) for 2 hours with rotation at 4°C. After three washes with cold DPBS, Sepharose coupled with protein G was added to the protein lysate-antibody mixture and incubated for 90 minutes at 4°C under rotation. The protein-antibody-sepharose mixture was then washed 5 times in Pierce IP lysis buffer (ThermoFisher Scientific) and resuspended in 2X Laemmli sample buffer (Sigma). Samples were then incubated at 95°C for 5 minutes and stored at -80°C until western blot analysis.

### Immunofluorescence

Cells were plated on Geltrex-coated glass coverslips. Cultured cells were fixed in 4% paraformaldehyde (PFA) in PBS (Alfa Aesar) for 10 minutes at r.t. followed by two washes in DPBS.

Human fetal tissue from terminated pregnancies was obtained from the joint MRC-Wellcome Trust Human Developmental Biology Resource (HDBR, http://hdbr.org, (Gerrelli et al., 2015)) which has been granted approval to function as a Research Tissue Bank (by the National Research Ethics Service (NRES) under research ethics committee approvals 18/LO/0822 and Newcastle 18/NE/0290). For immunostaining experiments, the fetal brains were fixed with 4% PFA in PBS (Alfa Aesar) for at least 24 hours at 4°C. After fixation, brains were dehydrated in graded ethanol washes and embedded in paraffin, before being cut and mounted on slides (the Experimental Histopathology Laboratory, the Francis Crick Institute).

Both cells and tissue were subjected to antigen retrieval in 10mM Na citrate: boiled 10 minutes at 95°C for adherent cells, and 30 seconds in the microwave for brain tissue. Samples were permeabilized in 0.1% Triton-PBS for 10 minutes at r.t. with rocking, blocked with 10% normal donkey serum (NDS, Jackson ImmunoResearch) in 0.1% Triton-PBS for 1 hour at r.t. with rocking, and subsequently incubated in primary antibodies (Supplemental Table 3) diluted in 10% NDS in 0.1% Triton-PBS overnight at 4°C with rocking. The next day, samples were washed 3 times in PBS, and incubated in secondary antibodies (Supplemental Table 3) and DAPI (10g/ml, Sigma,) diluted in 10% NDS in 0.1% Triton-PBS for 90 minutes at r.t. with rocking. Following 3 washes in PBS, samples were mounted in Vectashield mounting medium (Vector Laboratories). Immunofluorescence was performed on a minimum of three biological replicates from independent *in vitro* neuronal differentiations.

### Proximity Ligation Assay

Proximity Ligation Assay (PLA) was performed using the Duolink In Situ Red Started Kit Mouse/Rabbit (Sigma Aldrich) according to manufacturer’s instructions. Briefly, cells or human fetal brain tissue were subjected to fixation, antigen retrieval, and permeabilization as described above. Samples were then blocked in Duolink Blocking Solution for 60 minutes at 37°C, followed by incubation in primary antibodies (Supplemental Table 3) diluted in Duolink Antibody Diluent overnight at 4°C. The next day, samples were washed two times in Duolink Wash Buffer A, followed by incubation with PLA PLUS and MINUS probes for 1 hour at 37°C, ligation using the Duolink ligase diluted in the 5X Duolink Ligation Buffer for 30 minutes at 37°C and amplification using Duolink polymerase diluted in the 5X Duolink Amplification Buffer for 100 minutes at 37°C. All samples were washed twice in Duolink Wash Buffer A and once in Duolink Wash Buffer B, followed by mounting in Duolink In Situ Mounting Medium with DAPI.

### Image acquisition

Imaging was performed using a laser scanning TCS SP5 II confocal microscope (Leica Microsystems) at a z-section thickness of 1µm. The same settings were applied to all images. Images were visualized with FIJI (Schindelin et al., 2012)). Analysis of images for PLA was carried out in FIJI (Schindelin et al., 2012)): Maximum-intensity Z projection was performed followed by nuclei segmentation using Stardist (Schmidt et al., 2018), and dots detected via Fiji’s Find Maxima function. Dots occurring within segmented nuclei were then counted automatically on a per-nucleus basis.

### Intracellular flow cytometry

24hrs after exposure to the NOTCH inhibitor DAPT (Tocris), cells were washed in DPBS and dissociated with Accutase (Sigma) using a P1000 pipette. Once detached, cells were collected into 15 ml tubes with 4 volumes of DMEM-F12 (ThermoFisher Scientific) and pelleted at 300g, 3min. Cells were resuspended in DPBS, pelleted, then resuspended in Live/Dead™ Fixable Near-IR Dead Cell Stain (Invitrogen) as per manufacturer’s instructions, and left to incubate for 30mins, protected from light at r.t.. Cells were subsequently pelleted, and washed once in DPBS, before fixation in 4% paraformaldehyde in PBS. Following 10mins incubation, cells were washed with DPBS by dilution, then pelleted and resuspended in PBS before storage at 4°C for future analysis.

On the day of analysis, cells were transferred for staining into V-bottomed 96-well plates (Corning). Samples were pelleted (1000 x g, 4mins, 4°C) and resuspended in 100μl primary antibodies (Supplemental Table 3) diluted in PBS□+□0.2% Triton X-100 + 3% Donkey serum (Jackson), and incubated overnight at 4°C. The next day, samples were washed by dilution with 100μl PBS, pelleted and resuspended in 100μl PBS to complete the wash. After washing, cells were pelleted and resuspended and incubated in 100μl secondary antibodies (Supplemental Table 3) + 1μg/ml DAPI for 1hr. One additional wash was performed before transferring into cell strainer-capped tubes (Falcon) for acquisition on a Fortessa flow cytometer (BD) using FACSDiva software. Analysis was subsequently performed in FlowJo.

### RNA-seq and analysis

Libraries were prepared using the KAPA mRNA polyA HyperPrep kit (Illumina) and subsequently sequenced on the Illumina HiSeq4000 platform (Advanced Sequencing Facility, the Francis Crick Institute) to generate 100bp paired-end reads.

For sequence analysis, adapter trimming was performed with cutadapt (Martin, 2011) with parameters “--minimum-length=25 --quality-cutoff=20 -a AGATCGGAAGAGC”. The RSEM package (Li and Dewey, 2011) in conjunction with the STAR alignment algorithm (Dobin et al., 2013) was used for the mapping and subsequent gene-level counting of the sequenced reads with respect to the human UCSC hg19 genome and annotation (UCSC (Karolchik et al., 2004)) downloaded from AWS iGenomes (https://github.com/ewels/AWS-iGenomes). The parameters passed to the “rsem-calculate-expression” command were “--star --star-gzipped-read-file --star-output-genome-bam --forward-prob 0”. Differential expression analysis was performed with the DESeq2 package (Love et al., 2014) within the R programming environment (v3.3.1). An adjusted p-value of <= 0.05 and a fold change >= 1.5 was used as the significance threshold for the identification of differentially expressed genes.

For gene ontology enrichment the online functional annotation tool of the DAVID version 2021 bioinformatics resource https://david.ncifcrf.gov/summary.jsp was used. A background gene list was generated from genes expressed in wild type DIV24 cultures where RSEM computed expected count was ≥10 in at least one replicate. The “GOTERM_BP_DIRECT”, “GOTERM_CC_DIRECT” and “GOTERM_MF_DIRECT” functional annotation terms were selected to identify statistically enriched gene ontology annotations within sets of gene IDs associated with differentially expressed transcripts, and to calculate associated Benjamini–Hochberg adjusted *p* values.

### ChIP-seq

The ChIP-seq protocol was modified from (Sullivan and Santos, 2020). Cells were fixed with 2nM di(N-succimidyl) glutarate (Sigma-Aldrich) in DPBS (ThermoFisher Scientific) for 45 minutes at r.t. on a rocking platform. Three DPBS washes were then performed, before a second 10-minute fixation in 1% methanol-free formaldehyde solution (ThermoFisher Scientific) in DPBS at r.t. with rocking. The formaldehyde fixation was stopped by adding 1ml of 1.25M glycine (Sigma-Aldrich), followed by a 5-minute incubation on the rocking platform at r.t.. Cells were scraped off and pelleted by centrifugation at 800 x g for 5 minutes at 4°C. After three washes with ice-cold DPBS, the cell pellet was snap frozen in liquid nitrogen and stored at -80°C until processing. To isolate nuclei, pellets were resuspended in 300ml of SDS Lysis Buffer (Supplemental Table 1) containing 1X Protease inhibitor cocktail (ThermoFisher Scientific) and incubated for 30 minutes on ice. The cell suspension was transferred to a 1.5ml Diagenode TGX tube (Diagenode) and sonicated for 75 cycles (30 seconds on, 30 seconds off, on High) in a precooled Diagenode Bioruptor Plus Sonication System. 1 ml of Chromatin Dilution buffer (Supplemental Table 1) containing 1X Protease inhibitor cocktail was added to the crosslinked sheared chromatin and centrifuged at 14,000 x g for 30 minutes at 4°C. 60ml of soluble chromatin were stored at -20° as input chromatin, while the remaining supernatant was transferred to a protein LoBind tube (Fisher Scientific) containing protein A or G dynabeads (ThermoFisher Scientific) – primary antibody (Supplemental Table 3) mix (previously incubated with rotation for 3 hours at r.t.). The chromatin-antibody-dynabeads solution was incubated with rotation at 4°C overnight. Using a magnetic holder to separate the dynabeads, the supernatant was removed and sequentially washed for 5 minutes at 4°C under rotation with 1ml of Wash buffer A, Wash buffer B, Wash buffer C, twice with TE buffer (Supplemental Table 1). 100ml of Elution buffer (Supplemental Table 2) was added after the final wash, followed by a 5-minute incubation at 65°C. Dynabeads were separated using a magnetic holder and the eluted DNA was transferred to a clean 1.5ml tube. The elution step was repeated, resulting in 200ml of final DNA. The input chromatin from day 1 was removed from the freezer and the NaCl (Sigma-Aldrich) concentration was increased to 160mM for all samples. RNase A (ThermoFisher Scientific) was added to a final concentration of 20mg/ml and all samples were incubated at 65°C overnight to reverse crosslinks and digest contaminating RNA. On day 3, EDTA concentration was increased for all samples to 5mM (Sigma-Aldrich) followed by a 2-hour incubation with 200ug/ml Proteinase K (Sigma-Aldrich) to digest proteins.

ChIP and input samples were purified using the Zymo Clean and Concentrator-5 kit (Zymo) according to manufacturer’s instructions. DNA fragment size and distribution was determined by Agilent TapeStation (Agilent, USA) before DNA library preparation using the NEB Ultra II DNA Library Prep Kit for Illumina (New England BioLabs) as per manufacturer’s instructions. ChIP-seq samples were sequenced on the Illumina HiSeq4000 platform (Advanced Sequencing Facility, the Francis Crick Institute) and 100bp single-end reads were generated.

### ChIP-seq analysis

The nf-core/ChIP-seq pipeline (Ewels et al., 2020; https://doi.org/10.5281/zenodo.3529400) written in the Nextflow domain specific language (Di Tommaso et al., 2017) was used to perform the primary analysis of the samples in conjunction with Singularity (Kurtzer et al., 2017). The command used was “nextflow run nf-core/chipseq --input design.csv --genome hg19 min_reps_consensus 2 -profile crick -r 1.1.0”; where applicable the “--single_end” and “--narrow_peak” (for ASCL1 immunoprecipitations) or “--broad_peak” (for SMARCB1, H3K4me3 and H3K27ac immunoprecipitations) parameters were used. To summarize, the pipeline performs adapter trimming (Trim Galore! - https://www.bioinformatics.babraham.ac.uk/projects/trim_galore/), read alignment (BWA (Li and Durbin, 2009)) and filtering (SAMtools (Li et al., 2009); BEDTools (Quinlan and Hall, 2010); BamTools (Barnett et al., 2011); pysam - https://github.com/pysam-developers/pysam; picard-tools - http://broadinstitute.github.io/picard), normalized coverage track generation (BEDTools (Quinlan and Hall, 2010); bedGraphToBigWig (Kent et al., 2010)), peak calling (MACS (Zhang et al., 2008)) and annotation relative to gene features (HOMER (Heinz et al., 2010)), consensus peak set creation (BEDTools (Quinlan and Hall, 2010)), differential binding analysis (featureCounts (Liao et al., 2014); R - R Core Team (2017); DESeq2 (Love et al., 2014)) and extensive QC and version reporting (MultiQC (Ewels et al., 2016); FastQC - https://www.bioinformatics.babraham.ac.uk/projects/fastqc/; preseq (Daley and Smith, 2013); deepTools (Ramírez et al., 2016); phantompeakqualtools (Landt et al., 2012)). All data was processed relative to the human UCSC hg19 genome build (UCSC (Karolchik et al., 2004)) downloaded from AWS iGenomes (https://github.com/ewels/AWS-iGenomes). Peak annotation was performed relative to the same GTF gene annotation file used for the RNA-seq analysis. Tracks illustrating representative peaks were visualized using the IGV genome browser (Robinson et al., 2011).

Motif enrichment analyses of ChIP-Seq peak datasets were performed using HOMER (Heinz et al., 2010) findMotifsGenome with default parameters and region size set to 200bp bp (±100 bp adjacent to peak center): “findMotifsGenome.pl <peak/BED file> <genome> <output directory> - size 200”.

### ATAC-seq

ATAC-seq sample preparation was performed using previously established protocols (Buenrostro et al., 2013; Buenrostro et al., 2015; Corces et al., 2017). Briefly, 50,000 cells at each stage were isolated and pelleted at 500 x g for 5 minutes at 4°C, lysed in 50ml of ice-cold ATAC Resuspension Buffer (RSB, Supplemental Table 1) containing 0.1% NP40 (Sigma-Aldrich), 0.1% Tween-20 (Sigma-Aldrich) and 0.01% Digitonin (Promega) and incubated on ice for 3 minutes. The lysis reaction was stopped with 1ml of ice-cold ATAC-RSB containing 0.1% Tween-20 (Sigma-Aldrich) and the nuclei extracts isolated by centrifugation at 500 x g for 10 minutes at 4°C. The cell pellet was then re-suspended in 50ml of transposition reaction mix (25ml 2x TD buffer (Illumina), 2.5ml transposase (Illumina), 16.5ml DPBS (ThermoFisher Scientific), 0.5ml 1% digitonin (Promega), 0.5ml 10% Tween-20 (Sigma-Aldrich), 5ml water (ThermoFisher Scientific)) and subsequently incubated at 37°C for 30 minutes in a thermomixer with 1000 RPM mixing. Transposed DNA was purified using the Zymo Clean and Concentrator-5 kit (Zymo) according to the manufacturer’s protocol and eluted in 21ml of elution buffer. 5ml of the cleaned transposed DNA was used for library amplification (12 cycles) using the NEBNext HiFi 2X PCR Master Mix (New England BioLabs) and previously designed ATAC-seq barcoded primers (Supplemental Table 4) (Buenrostro et al., 2013). PCR reactions were cleaned-up with KAPA pure beads (Roche) at 1.8X beads versus sample ratio. Prior to sequencing, DNA fragment size and distribution and library concentration were determined by Agilent TapeStation (Agilent, USA) and QubitTM dsDNA HS assay (Life Technologies), respectively. ATAC-seq samples were subsequently sequenced on the Illumina HiSeq4000 platform (Advanced Sequencing Facility, the Francis Crick Institute) and 100bp paired-end reads were generated.

### ATAC-seq analysis

The nf-core/atacseq pipeline ((Ewels et al., 2020); https://doi.org/10.5281/zenodo.3529420) written in the Nextflow domain specific language (Di Tommaso et al., 2017) was used to perform the primary analysis of the samples in conjunction with Singularity (Kurtzer et al., 2017). The command used was “nextflow run nf-core/atacseq --design design.csv –genome hg19 - min_reps_consensus 1 -profile crick -r 1.1.0”. The nf-core/atacseq pipeline uses similar processing steps as described for the nf-core/chipseq pipeline in the previous section but with additional steps specific to ATAC-seq analysis such as the removal of mitochondrial reads.

For peak intersection, BEDtools intersectBed was used to identify genomic intervals overlapping by 1bp in BED files listing coordinates of consensus peak sets: “bedtools intersect -a file1.bed -b file2.bed > output.txt”.

### Single cell RNA-seq and analysis

On the day of harvest, adherent cell cultures cells were washed twice with DPBS then incubated at 37°C in accutase (Sigma-Aldrich) and manually dissociated by pipetting with a P1000. 2 volumes of DMEM F-12 were added and cells passed through a 40µm cell strainer to achieve a single-celled suspension before counting on a NucleoCounter NC-200. 1.5 × 10^6^ cells were then centrifuged at 255 x g for 2mins. The pellet was resuspended in 750ml of 0.22µm-filtered 1% BSA in DPBS for a theoretical concentration of 2000 cells/ml. This suspension was then counted. 10,000 cells were loaded onto the Chromium X (10x Genomics) to generate single-celled Gel Beads-in-Emulsion (GEMs) for library preparation. Dual index Chromium Single Cell 3’ v3.1 Chemistry was used according to manufacturer’s instructions, using 11 cycles for the cDNA amplification and 14 cycles for the library PCR. Libraries were then sequenced on a NovaSeq 6000 (Illumina).

#### Pre-processing

Feature quantification of the spliced and unspliced reads was performed using alevin (Srivastava et al., 2019) as recommended by Soneson et al. 2021. Reads mapped to the human reference genome (ensembl GRCh37, release 75). The quantification was carried out in two stages. In the first pass, Cell Ranger Single-Cell Software Suite from 10X Genomics was availed to identify filtered whitelist of cell barcodes. This whitelist of cell barcodes was used as input for alevin quantification in second pass. This helped use the Cell Ranger’s default filtering criteria for selection of cell barcodes for downstream analysis. Further analysis was carried out using Seurat package (Butler et al., 2018; Hao et al., 2021; Stuart et al., 2019) in R-4.0.0 (R Core Team, 2020). A sample specific threshold for low-quality cells was identified using median absolute deviation (MAD) measures for cells expressing > 3 MAD of mitochondrial gene expression along with < 3 MAD of total number of detected features/genes. Suspected doublet cells were identified using scDblFinder v1.4.0 (Germain et al., 2021) and were excluded from subsequent analysis. Using default parameters within Seurat package (Butler et al. 2018, Stuart et al. 2019, Hao et al. Cell 2021) each sample was normalized and the variance stabilized across cells using the SCTransform v0.3.3 (Hafemeister and Satija, 2019) with glmGamPoi (Ahlmann-Eltze and Huber, 2021) method.

#### Integration across samples

After filtering of cells/clusters based on mitochondrial gene expression and the number of detected features, we performed following sets of integration of the samples using the standard workflow from the Seurat package (Butler et al., 2018; Hao et al., 2021; Stuart et al., 2019). These include integration of samples from individual sample groups (WT cells with and without DAPT treatment and *ASCL1* KO cells with DAPT treatment) and by DAPT treatment or genotype (a) WT cells with and without DAPT treatment (Figure S1B) and b) WT cells and *ASCL1* KO cells with DAPT treatment (Figure S2C)). After data normalization and variable feature detection in the individual samples using SCTransform (Hafemeister and Satija, 2019) (see above), anchors were identified using the ‘FindIntegrationAnchors()’ function and datasets were integrated with the ‘IntegrateData()’ across 30 dimensions for 3000 integration features identified from the datasets. We then performed PCA on the integrated data and the first 20 principal components were used to create a Shared Nearest Neighbor (SNN) graph using the ‘FindNeighbors()’ function. This was used to find clusters of cells showing similar expression using the ‘FindClusters()’ function across a range of clustering resolutions (0.2-1.4 in 0.2 increments). For data visualization dimensional reduction of the integrated data was achieved using UMAP with 20 principal components and with cosine correlation metric. For further downstream analysis, the raw RNA counts of the integrated data were used after normalization with the “LogNormalize” method.

For the integrated dataset of WT cells treated with DAPT, clustering resolution of 1.0 (Figure 1B) was selected based on visual inspection of known marker genes (Figure 1F, gene list available at Supplemental Table 5) that are enriched in individual clusters. Cell identity was determined based on the module score identified for these marker genes using ‘AddModuleScore()’ function. Using this integrated data as a reference, cell identities for the ASCL1 KO dataset were identified with the help of transfer anchors obtained using the first 20 principal components of SCTransform (Hafemeister and Satija, 2019) normalized features (Figure 3A).

For the integrated datasets of a) all WT cells with or without DAPT (Figure S1B), and b) WT and ASCL1 KO cells with DAPT treatment (Figure S2C), cell identities were determined by visual inspection based on their module scores of the same marker genes, like above.

#### RNA velocity estimation

The spliced and unspliced read counts from the integrated Seurat object was used as input into scVelo (Bergen et al., 2020) to calculate RNA velocity values for each gene of each cell. scVelo was used in the “dynamical” mode with default settings. The resulting RNA velocity vector was embedded into the PCA and UMAP space by translating the RNA velocities into likely cell transitions using cosine correlation to compute the probabilities of one cell transitioning into another cell. We identified driver genes, i.e., those genes that show dynamic behavior, as those genes with a fit likelihood in the dynamical model > 0.3. We also used PAGA (Wolf et al., 2019) to perform trajectory inference for which directionality was inferred from the RNA velocities.

### IGV visualisation

ChIP-seq and ATAC-seq datasets were visually explored using the Interactive Genomics Viewer (IGV) (Robinson et al., 2011; Thorvaldsdóttir et al., 2013). Images of genomic regions used as proof of principle to demonstrate binding and/or accessibility dynamics within different genotypic conditions were exported as .svg files and cropped with Adobe Illustrator (Adobe, USA).

### Global data visualisation

All global visualization of specific ChIP-seq and ATAC-seq datasets (heat maps) was performed using deepTools (Ramírez et al., 2016). The combination of the following two commands was used to generate coverage heatmaps that display an average of normalized read density for specific subsets of peaks: computeMatrix reference − point −S (.bigWig) <> −R <.bed> <> n −o <files.gz> −b 1000/3000 −a 1000/3000 −−referencePoint center and plotHeatmap −m <f i l e s. gz> −o <Heatmap.svg> n −−colorMap # −−heatmapHeight # −−heatmapWidth #

### Activity-by-contact (ABC) algorithm

Enhancer-gene connections were established using the Activity-by-Contact (ABC) model (Fulco et al., 2019) for the wildtype condition using the information obtained from the ATAC-seq (MACS “–narrow_peak” calling mode), the H3K27ac ChIP-seq (MACS “–broad” peak calling mode) and RNA-seq in wildtype DIV24 cultures. Each datatype was analysed as specified above. ABC scores for each gene and chromatin accessible element within a 5Mb range were calculated. To generate the neccessary gene and TSS annotation files we used the GRCh37.75 annotation in R-3.6.2 (R Core Team, 2019) and the Bioconductor package plyranges (version 1.14.0,(Lee et al., 2019)). Transcription start sites (TSS) for each gene were selected based on the most highly expressed isoform (highest mean TPM expression across the three replicates in the RNA-seq). In cases in which several isoforms show equal expression levels, we selected the TSS that is used by the majority of isoforms. Lastly, for the remaining genes, i.e. those for which neither gene expression nor the majority vote identified a unique TSS, we selected the most 5’ TSS. The TSS region was then defined as the 500bp surrounding each gene’s TSS. We removed genes corresponding to small RNAs (gene symbol contains “MIR” or “RNU”, genes with a gene body length < 300bp (we calculated the gene body length by summing across the exon widths of each transcript)). For the gene annotation each gene was collapsed its most expanded genomic ranges.

#### Define candidate elements

Instead of the makeCandidateRegions.py script we used the Bioconductor package DiffBind (Ross-Innes et al., 2012). We run MACS (Zhang et al., 2008) for each ATAC-seq replicate using the ABC algorithm-specific parameters (-p 0.1 --call-summits TRUE) and removed elements overlapping regions of the genome that have been observed to accumulate anomalous number of reads in epigenetic sequencing available via the ENCODE project (ENCODE Project Consortium (Luo et al., 2019)) for GRCh37 with the following identifier ENCSR636HFF. Subsequently, reads were counted with DiffBind::dba.count(DBA, summits = 275, minOverlap=2) in the consensus peaks identified with DiffBind. Peaks in the consensus peak set were re-centred and trimmed based on their points of greatest read overlap (summits) to provide more standardized peak intervals. After background normalisation, candidate putative enhancer regions were identified as those 150000 consensus peaks with the highest mean normalized read count. Finally, we merged the candidate putative enhancer regions with the annotated TSS file region (“include-list”), as the ABC model considers promoters as part of the putative enhancer set.

#### Quantifying enhancer activity

The activity of the putative enhancer regions was then quantified using the run.neighborhoods.py function from the ABC algorithm including the information for the RNA-sequencing to define expressed genes.

#### Computing the ABC score

Finally, ABC scores were calculated using the predict.py without experimental contact data information (using the following parameters: –score_column powerlaw.Score –threshold .022 – make_all_putative).

### Statistics

Where data are presented as the mean ± standard error of the mean (SEM), unpaired Student’s t-test was used to determine statistical significance (GraphPad Prism and R). Details of statistical analyses are found in the figure legends.

## Acknowledgments

We gratefully acknowledge the Human Embryo Stem Cell Unit, Advanced Sequencing Facility, Flow Cytometry, Advanced Light Microscopy and Proteomics facilities at the Francis Crick Institute. We thank Adrienne Sullivan from the Quantitative Cell Biology lab, and Dinis Calado from the Immunity and Cancer Lab, for experimental advice and members of the Neural Stem Cell Biology lab for feedback on the manuscript. This work was supported by the Francis Crick Institute, which receives its funding from Cancer Research UK (FC0010089), the UK Medical Research Council (FC0010089), and the Wellcome Trust (FC0010089). This work was also supported by the Wellcome Trust (Career Development Fellowship 209568/Z/17/Z to C.D., and Investigator Award 106187/Z/14/Z to F.G.).

## Author contributions

Conceptualization: O.P., C.D. and F.G.; Methodology: O.P., H.P., S.S., Y.X.T., S-L.A., C.D. and F.G.; Software: S.S., A.G.; Validation: O.P.; Formal Analysis: O.P., S.S., A.G., Y.X.T., C.D.; Investigation: O.P., Y.X.T., H.P., S.S., A.G., C.C.G., S-L.A., M.L.S., L.G., C.D.; Resources: S-L.A.; Data Curation: O.P., Y.X.T., S.S., A.G., C.D.; Writing – Original Draft: O.P., F.G. and C.D.; Writing – Review & Editing: all authors; Visualization: O.P., Y.X.T., A.G.; Supervision: F.G. and C.D.; Funding Acquisition: F.G.; Project Administration: F.G. and C.D..

## Declaration of interests

The authors declare no competing interests.

## Data availability

scRNA-Seq, RNA-seq, ATAC-seq and ChIP-seq data generated in this study are available for download at GSE214383.

## Supplemental information

**Figure S1**. Related to Figure 1. ASCL1 expression marks a transitional cell population bridging actively dividing progenitors and postmitotic neurons.

**Figure S2**. Related to Figure 3. Generation of Transitional Progenitors and Neurons is impaired in *ASCL1* KO DIV24 cultures.

**Figure S3**. Related to Figure 5. mSWI/SNF npBAF and nBAF subunits in human iPSC-derived neural cultures.

**Figure S4**. Related to Figures 4 and 6. Interference with mSWI/SNF ATPase activity confirms co-dependency of ASCL1 and mSWI/SNF at a subset of regulatory elements

**Figure S5**. Related to Methods. Differential transcriptomic analysis of DIV24 neuronal cultures derived from three different iPSC lines.

**File S1**. Differential gene expression analysis of *ASCL1* KO vs control neural cultures at DIV24.

**File S1**. Distal regulatory element map in DIV24 wild type neural cultures predicted by the ABC algorithm (annotated to GRCh37).

**File S3**. Related to Figure 3E,F. ASCL1-regulated genes: differential gene expression and gene ontology analysis.

**File S4**. Motifs from HOMER known motif analysis.s

**Supplemental Table 1**. Media and buffer recipes.

**Supplemental Table 2**. List of qRT-PCR TaqMan probes.

**Supplemental Table 4**. ATAC-seq primers with barcodes (Buenrostro et al., 2013).

**Supplemental Table 5**. List of genes used to group DIV24 cells into three populations: cycling progenitors, transitional progenitors, and neurons.

## Notes

### Competing Interest Statement

The authors have declared no competing interest.

