## Supplemental Figure for "ASCL1 interacts with the mSWI/SNF at distal regulatory elements to regulate neural differentiation"

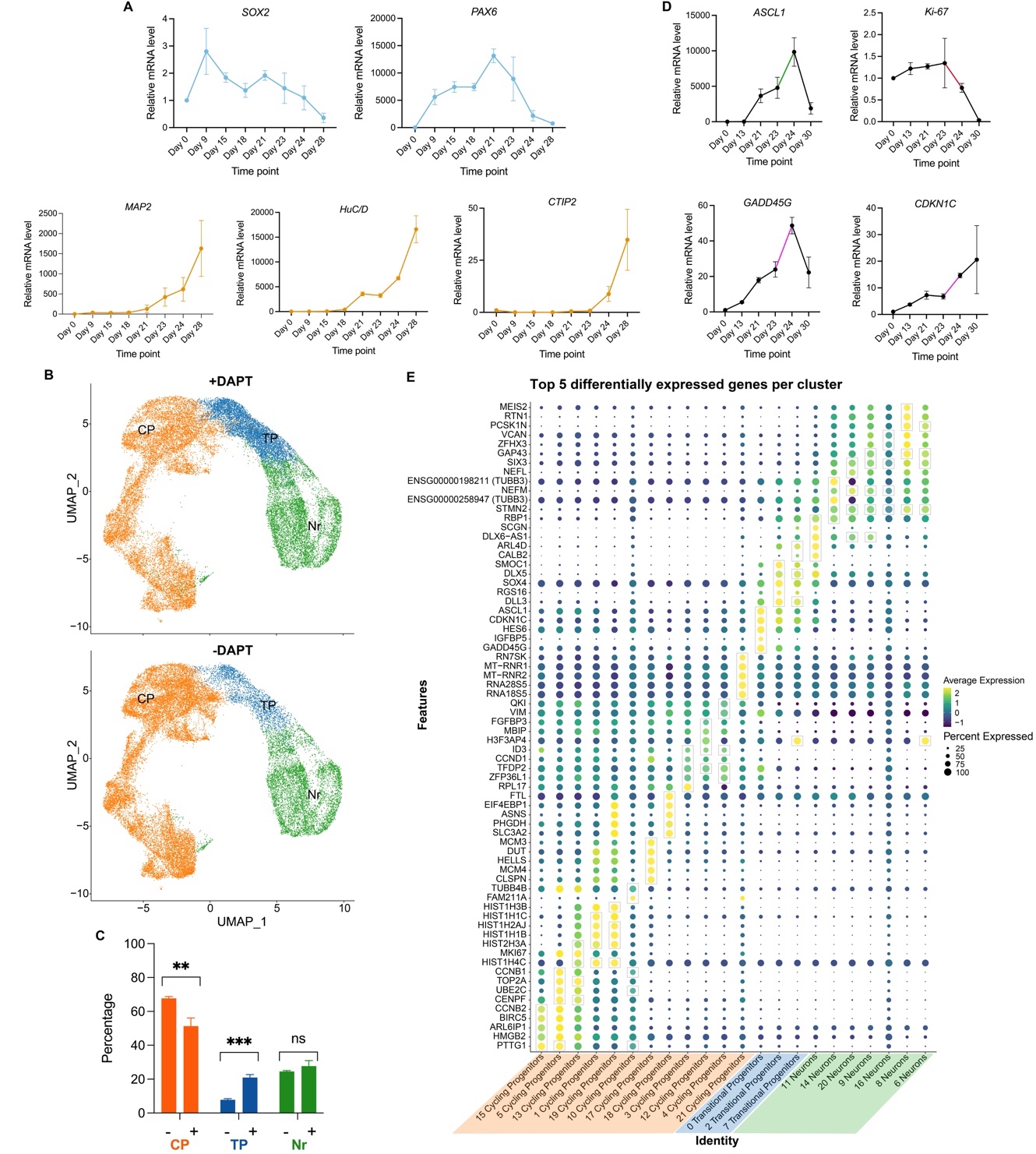

### Figure S1, related to Figure 1. ASCL1 expression marks a transitional cell population bridging actively dividing progenitors and postmitotic neurons.

**(A)** qRT-PCR analysis of the expression of neural progenitor-associated genes *SOX2* and *PAX6* (top) activated after neural induction with dual SMAD inhibition; mRNA expression relative to DIV0. Onset of expression of neuronal genes *MAP2*, *HUC/D* (*ELAVL3/4*) and *CTIP2* (*BCL11B*) is between DIV18 and 21, with marked increase from DIV24 (post Notch inhibition with DAPT) (bottom). Data from same experiment as Figure 1A (three independent cultures). Error bars represent mean ±SEM for three biological replicates. **(B)** UMAP plot projections of single-cell transcriptomes of DIV24 neural cultures, collected from cultures both with and without Notch inhibition at DIV23; a UMAP embedding was estimated after integration of datasets (three replicates in each condition) and then cell types were assigned using the markers shown in Figure 1F; different experiments with (+DAPT) and without (-DAPT) Notch inhibition are shown on the integrated UMAP. Dots represent single cells. Colors represent the different clusters defined by marker expression. **(C)** Relative proportion of different cell state clusters from each dataset from (B) showing significant enrichment of Transitional Progenitors with DAPT addition. Unpaired t-test, **p<0.01, ***p<0.001. **(D)** qRT-PCR analysis of the expression of *ASCL1*, *MKI67* (cell proliferation marker) and *CDKN1C* and *GADD45G* (cell cycle exit markers) at multiple timepoints during neural differentiation; mRNA expression relative to DIV0. Colored lines highlight expression changes upon addition of Notch inhibitor DAPT: the increase in *ASCL1* expression between DIV23 and DIV24 is accompanied by a decrease in *MKI67* and increase in *GADD45G* AND *CDKN1C*. Data obtained from an independent experiment from Figure 1A and panel (A). Error bars represent mean ±SEM for three biological replicates. **(E)** Dot plot representation of the expression of all the top 5 genes enriched in the 22 clusters identified by Seurat in Figure 1C (the top 5 for each cluster are outlined in boxes) in comparison to the rest of the cells in the dataset. Dot size indicates percentage of cells in each cluster expressing a gene, shading indicates the average gene expression.

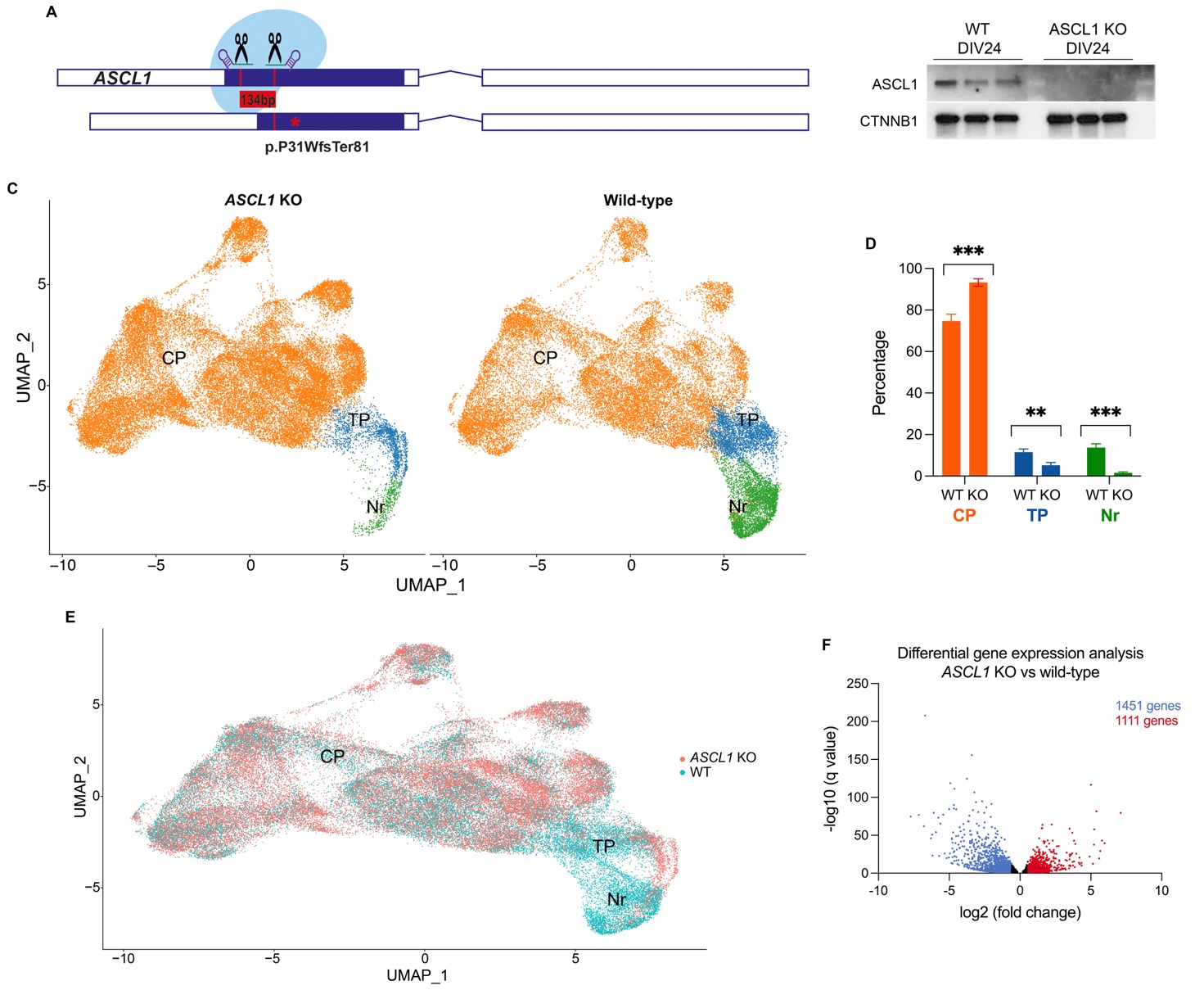

### Figure S2, related to Figure 3. Generation of Transitional Progenitors and Neurons is impaired in *ASCL1* KO DIV24 cultures.

**(A)** Diagram illustrating the 133bp deletion in the first exon of *ASCL1* generated by CRISPR/Cas9 (gRNA site targeting represented by scissors), inducing a frameshift with a premature stop codon (represented by a red asterisk), expected to undergo nonsense mediated decay. **(B)** Western blot analysis of ASCL1 protein expression in DIV24 neural cultures of three control (wild-type) and three *ASCL1* KO clones showing no protein expression (after 10 minutes exposure to chemiluminescent reagent). CTNNB1 loading control is included. **(C)** UMAP representations illustrating the three main cell populations (Cycling Progenitors, Transitional Progenitors and Neurons). *ASCL1* KO cells (left) and wild-type cells (right) are shown as UMAP plot for the integrated datasets from both conditions. **(D)** Proportion of cells in each of the three main populations (CP, Cycling Progenitors; TP, Transitional Progenitors; Nr, Neurons) in wild-type and *ASCL1* KO DIV24 neural cultures (derived from (C)). Unpaired t-test; ***p*<0.01, ****p*<0.001. **(E)** UMAP from (C) with cells grouped by genotype, showing that wild-type and *ASCL1* KO transitional progenitors and neurons cluster separately, suggesting transcriptome-wide differences in those cell types. **(F)** Volcano plot showing the genes dysregulated in *ASCL1* KO versus wild-type DIV24 cultures (bulk RNA-seq); genes with significant (fold change > 1.5, q-value < 0.05) differential gene expression are colored blue (downregulated) and red (upregulated).

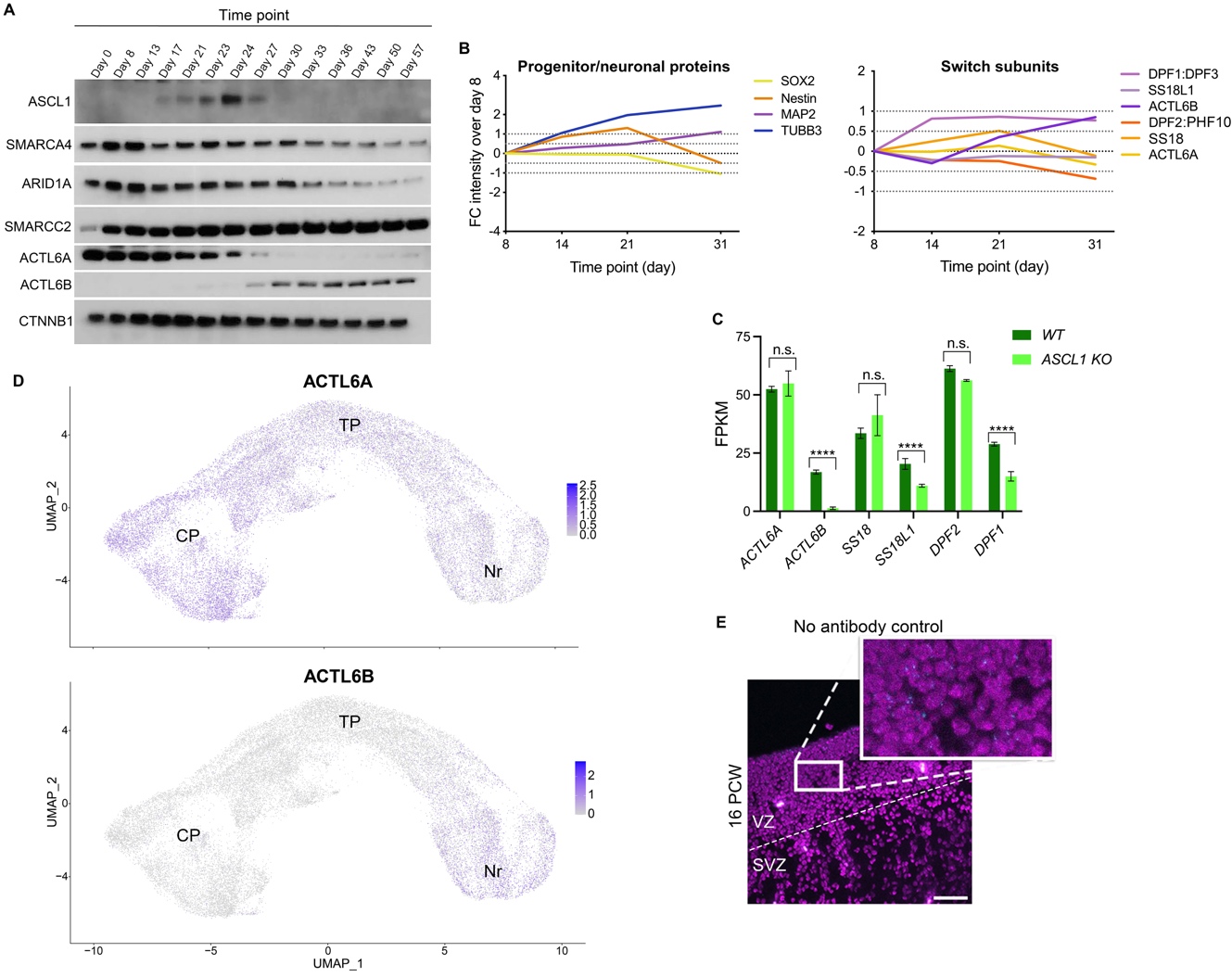

### Figure S3, related to Figure 5. mSWI/SNF npBAF and nBAF subunits in human iPSC-derived neural cultures.

**(A)** Western blot showing the expression patterns of ASCL1, mSWI/SNF core and switching subunits. CTNNB1 loading control is included. **(B)** Time course plots of proteins detected using TMT-labelled LC-MS/MS quantitation showing the decrease in expression of npBAF subunits DPF2, PHF10, ACTL6A during differentiation of neural cultures, parallel to the decrease in expression of the progenitor-specific markers SOX2 and Nestin. Conversely, nBAF subunits DPF1/DPF3, SS18L1, ACTL6B increase in expression, in parallel with the expression of the neuronal-specific markers MAP2 and TUBB3. **(C)** Normalized FPKM from RNA-Seq analysis for npBAF and nBAF subunits in wild-type and *ASCL1* KO DIV24 cultures. There is no change in the expression of npBAF subunits *ACTL6A*, *SS18*, or *DPF2*, while all nBAF subunits (*ACTL6B*, *SS18L1*, *DPF1*) are significantly downregulated. *****padj*<0.0001. **(D)** UMAP plots showing the expression of *ACTL6A* and *ACTL6B* in control cultures at DIV24; single cell gene expression is overlayed on the UMAP from Figure 1D. Transitional progenitors do not express nBAF subunit ACTL6B. **(E)** Representative immunofluorescence image of Proximity Ligation Assay no antibody control (related to Figure 5B) in the human fetal cortex at 16 PCW. Scale bar, 50um.

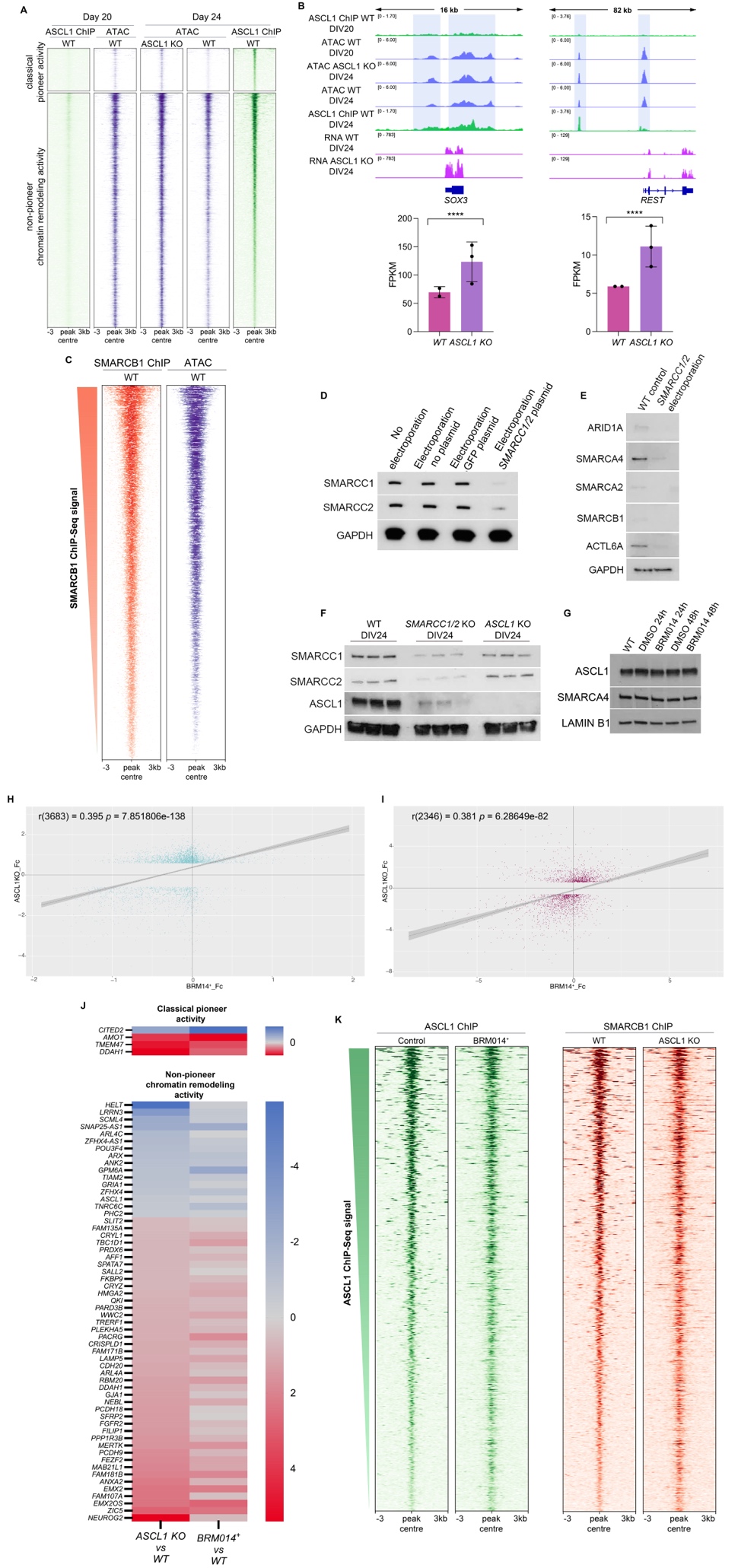

### Figure S4, Related to Figures 4 and 6. Interference with mSWI/SNF ATPase activity confirms co-dependency of ASCL1 and mSWI/SNF at a subset of regulatory elements

**(A)** Heatmaps representing the analysis of temporal dynamics of ASCL1 binding and chromatin accessibility that divides its pioneer activity into classical pioneer (top, n= 760) and non-pioneer chromatin remodeling (bottom, n= 4,108) activities at sites where ASCL1 represses accessibility. **(B)** Representative IGV tracks of relevant loci identified in (A). Bottom, bar plots show mean expression in FPKM for the depicted genes in wild-type and *ASCL1*-KO. *****padj* <0.0001. **(C)** Heatmaps representing SMARCB1 binding by ChIP-Seq (n=26,207) relative to chromatin state determined by ATAC-Seq in wild type DIV24 cultures. **(D)** Western blot for SMARCC1 and SMARCC2 on the collection day (DIV24) for cells that underwent electroporation with the plasmids targeting *SMARCC1* and *SMARCC2* and their respective controls. GAPDH loading control is included. **(E)** Western blot showing the protein levels of other mSWI/SNF subunits (ARID1A, SMARCA2, SMARCA4, SMARCB1, ACTL6A) in the SMARCC1/2 KO cells compared to control cells at DIV24. GAPDH loading control is included. **(F)** Western blot analysis of SMARCC1, SMARCC2 and ASCL1 expression in wild-type, *SMARCC1/2* KO and *ASCL1* KO DIV24 cultures. GAPDH loading control is included. **(G)** Western blot analysis of ASCL1 and SMARCA4 expression at 24 and 48h after treatment with ATPase inhibitor BRM014 (and vehicle control) showing treatment does not affect protein expression. LaminB1 loading control is included. **(H)** Scatterplot showing the change in accessibility assessed by ATAC-Seq in BRM014-treated cells versus control cultures at all ASCL1-bound sites that show a significant change in accessibility upon ASCL1 removal. Blue dots represent sites that also reach the significance threshold after 48hrs of BRM014 inhibitory treatment. Red dots represent sites that do not reach the significance threshold after 48hrs of BRM014 inhibitory treatment. Fc: fold change. **(I)** Scatterplot showing the changes at the transcriptional level assessed by RNA-Seq in BRM014-treated cells versus control cultures at all genes found to be significantly dysregulated upon ASCL1 removal. Blue dots represent genes that also reach the significance threshold after 48hrs of BRM014 inhibitory treatment (BRM sig). Fc: fold change. **(J)** Heatmaps showing the ABC-predicted genes dysregulated in both *ASCL1* KO and BRM014-treated cultures where ASCL1-SMARCB1 co-binding induces decreased accessibility at ASCL1 classical pioneer activity sites (top) and sites of ASCL1 non-pioneer chromatin remodeling activity (bottom). **(K)** Heatmaps profiling ASCL1 and SMARCB1 binding at ASCL1-mSWI/SNF dependent sites where the interaction is associated with closed chromatin, showing that interfering with mSWI/SNF ATPAse activity (BRM014 treatment) reduces ASCL1 binding and reciprocally, eliminating ASCL1 (*ASCL1* KO) reduces SMARCB1 binding.

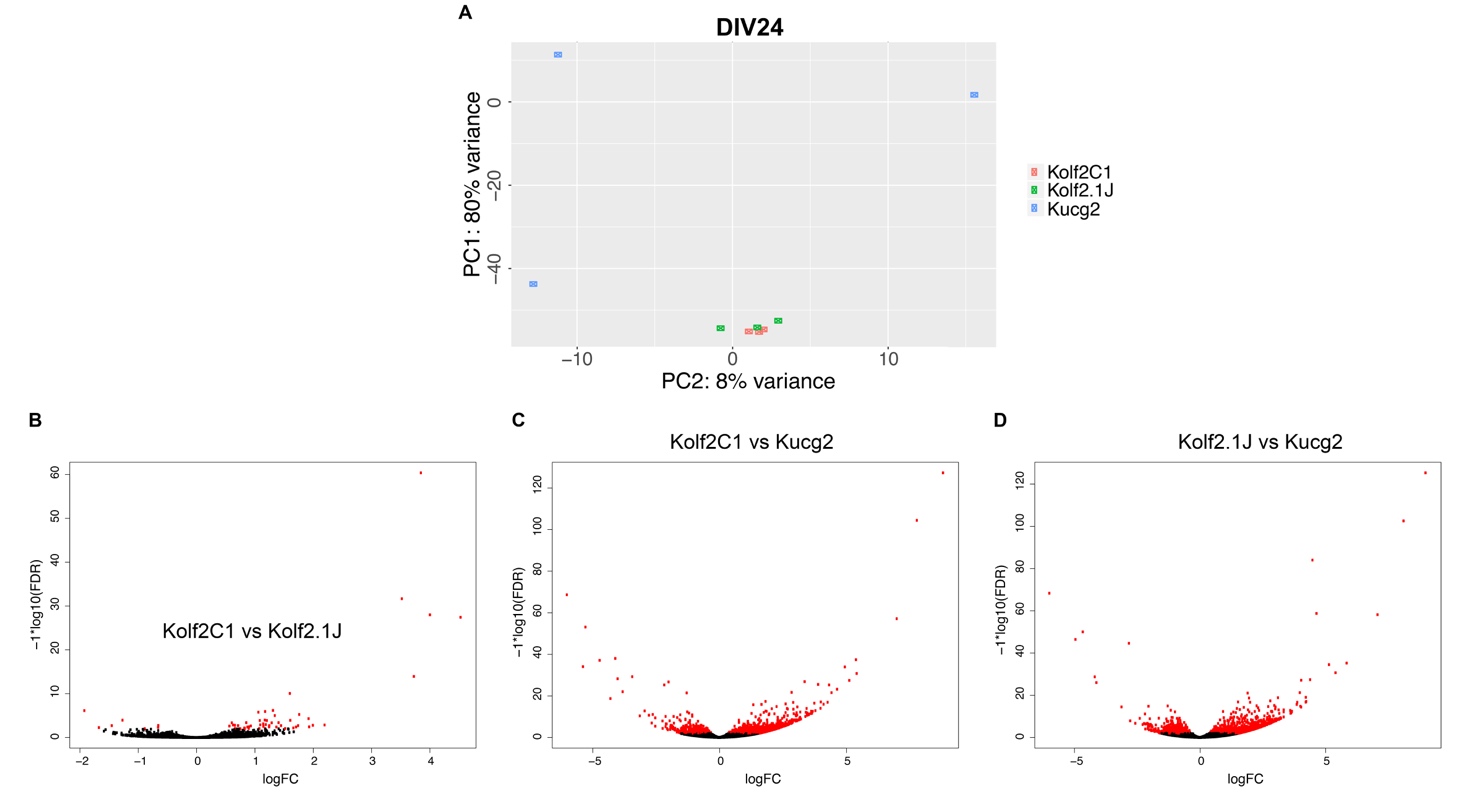

### Figure S5. Related to Methods. Differential transcriptomic analysis of DIV24 neuronal cultures derived from three different iPSC lines.

**(A)** PCA scatter plot of gene expression in three iPSC-line-derived neuronal cultures at DIV24: Kolf2C1, Kolf2.1J (CRISPR/Cas9 edited Kolf2C1) and independent line Kucg2. **(B, C, D)** Volcano plots representing differential gene expression analysis between pairs of the three lines tested in (A). Red dots indicate q<0.05.
