## Supplemental Table for "ASCL1 interacts with the mSWI/SNF at distal regulatory elements to regulate neural differentiation"

**Supplemental Table 1.** Media and Buffer Recipes

| **Component** | **Stock Concentration** | **Final Concentration** | **Volume** |
| --- | --- | --- | --- |
| **N2 media composition (500 ml)** | | | |
| DMEM/F-12 | - | 97% | 485 ml |
| N-2 Supplement | 100 X | 1 X | 5 ml |
| GlutaMAX-l Supplement | 100 X | 1 X | 5 ml |
| Penicillin/Streptomycin | 10,000 μg/ml (100X) | 100 μg/ml | 5 ml |
| **B27 media composition (500 ml)** | | | |
| Neurobasal Medium | - | 96% | 480 ml |
| B-27 Supplement | 50 X | 1 X | 10 ml |
| GlutaMAX-l Supplement | 100 X | 1% | 5 ml |
| Penicillin/Streptomycin | 10,000 μg/ml (100X) | 100 μg/ml | 5 ml |
| **ChIP SDS Lysis Buffer (50 ml)** | | | |
| Tris-HCl, pH 7.5 | 1M | 50mM | 2.5 ml |
| EDTA | 0.5M | 10mM | 1 ml |
| SDS | 20% | 1% | 2.5 ml |
| Water | - | - | 44 ml |
| **ChIP Chromatin Dilution Buffer (50 ml)** | | | |
| Tris, pH 7.5 | 1M | 25mM | 1.25 ml |
| EDTA | 0.5M | 5mM | 0.5 ml |
| Triton X-100 | 100% | 1% | 0.5 ml |
| SDS | 20% | 0.1% | 0.25 ml |
| Water | - | - | 47.5 ml |
| **ChIP Wash Buffer A (50 ml)** | | | |
| HEPES, pH 7.9 | 0.5M | 50mM | 5 ml |
| NaCl | 5M | 140mM | 1.4 ml |
| EDTA | 0.5M | 1mM | 0.1 ml |
| Triton X-100 | 100% | 1% | 0.5 ml |
| Sodium deoxycholate | 10% | 0.1% | 0.5 ml |
| SDS | 20% | 0.1% | 0.25 ml |
| Water | - | - | 42.25 ml |
| **ChIP Wash Buffer B (50 ml)** | | | |
| HEPES, pH 7.9 | 0.5M | 50mM | 5 ml |
| NaCl | 5M | 500mM | 5 ml |
| EDTA | 0.5M | 1mM | 0.1 ml |
| Triton X-100 | 100% | 1% | 0.5 ml |
| Sodium deoxycholate | 10% | 0.1% | 0.5 ml |
| SDS | 20% | 0.1% | 0.25 ml |
| Water | - | - | 38.65 ml |
| **Component** | **Stock Concentration** | **Final Concentration** | **Volume** |
| **ChIP Wash Buffer C (50 ml)** | | | |
| Tris, pH 8.0 | 1M | 20mM | 1 ml |
| EDTA | 0.5M | 1mM | 0.1 ml |
| LiCl | 8M | 250mM | 1.56 ml |
| NP-40 Alternative | 100% | 0.5% | 0.25 ml |
| Sodium deoxycholate | 10% | 0.5% | 2.5 ml |
| Water | - | - | 44.59 ml |
| **ChIP TE Buffer (50 ml)** | | | |
| Tris, pH 8.0 | 1M | 10mM | 0.5 ml |
| EDTA | 0.5M | 1mM | 0.1 ml |
| Water | - | - | 49.4 ml |
| **ChIP Elution Buffer (50 ml)** |  |  |  |
| Tris, pH 7.5 | 1M | 10mM | 0.5 ml |
| EDTA | 0.5M | 1mM | 0.1 ml |
| SDS | 20% | 1% | 2.5 ml |
| Water | - | - | 46.9 ml |
| **ATAC RSB Buffer (50 ml)** | | | |
| Tris-HCl, pH 7.4 | 1M | 10mM | 0.5 ml |
| NaCl | 5M | 10mM | 0.1mM |
| MgCl_2_ | 1M | 3mM | 0.15 ml |
| Water | - | - | 49.25 ml |

**Supplemental Table 2.** List of qRT-PCR TaqMan® probes

| **Target Gene** | **Supplier** | **Assay ID** | **Application** |
| --- | --- | --- | --- |
| *ASCL1* | ThermoFisher Scientific | Hs00269932_m1 | qRT-PCR |
| *BCL11B/CTIP2* | ThermoFisher Scientific | Hs01102259_m1 | qRT-PCR |
| *CDK1C* | ThermoFisher Scientific | Hs00175938_m1 | qRT-PCR |
| *GADD45G* | ThermoFisher Scientific | Hs00198672_m1 | qRT-PCR |
| *HPRT1* | ThermoFisher Scientific | Hs02800695_m1 | qRT-PCR |
| *HuC/D* | ThermoFisher Scientific | Hs00956610_mH | qRT-PCR |
| *MAP2* | ThermoFisher Scientific | Hs00258900_m1 | qRT-PCR |
| *MKI67* | ThermoFisher Scientific | Hs00606991_m1 | qRT-PCR |
| *PAX6* | ThermoFisher Scientific | Hs00240871_m1 | qRT-PCR |
| *SOX2* | ThermoFisher Scientific | Hs01053049_s1 | qRT-PCR |
| *UBC* | ThermoFisher Scientific | Hs00824723_m1 | qRT-PCR |

**Supplemental Table 3.** Resources Table

| **REAGENT or RESOURCE** | **SOURCE** | **IDENTIFIER** |
| --- | --- | --- |
| **Antibodies** | | |
| Rabbit monoclonal anti-ACTL6A | Abcam | Cat# ab131272;  RRID: [AB_11157110](http://antibodyregistry.org/AB_11157110) |
| Rabbit monoclonal anti-ACTL6B | Abcam | Cat# ab180927  RRID: AB_2924269 |
| Rabbit polyclonal anti-ARID1A | Bethyl | Cat# A301-041A  RRID: [AB_2060365](http://antibodyregistry.org/AB_2060365) |
| Mouse monoclonal anti-ASCL1 | BD Biosciences | Cat# 556604  RRID: [AB_396479](http://antibodyregistry.org/AB_396479) |
| Rabbit monoclonal anti-ASCL1 | Abcam | Cat# ab211327  RRID: AB_2924270 |
| Rabbit polyclonal anti-ASCL1 | Abcam | Cat# ab74065  RRID: [AB_1859937](http://antibodyregistry.org/AB_1859937) |
| Rat monoclonal anti-CTIP2 | Abcam | Cat# ab18465  RRID: [AB_2064130](http://antibodyregistry.org/AB_2064130) |
| Mouse monoclonal anti-GAPDH | Santa Cruz | Cat# sc-47724  RRID: [AB_627678](http://antibodyregistry.org/AB_627678) |
| Rabbit monoclonal anti-GAPDH | Cell Signaling Technology | Cat# 5174  RRID: [AB_10622025](http://antibodyregistry.org/AB_10622025) |
| Rabbit polyclonal anti-PAX6 | Covance | Cat# PRB-278P  RRID: [AB_291612](http://antibodyregistry.org/AB_291612) |
| Rabbit polyclonal anti-SMARCA4 | Santa Cruz | Cat# sc-10768  RRID: [AB_2255022](http://antibodyregistry.org/AB_2255022) |
| Mouse monoclonal anti-SMARCB1 | BD Biosciences | Cat# 612110  RRID: [AB_399481](http://antibodyregistry.org/AB_399481) |
| Rabbit polyclonal anti-SMARCB1 | Abcam | Cat# ab12167  RRID: [AB_298898](http://antibodyregistry.org/AB_298898) |
| Rabbit polyclonal anti-SMARCC1 | Abcam | Cat# ab72503  RRID: [AB_1270780](http://antibodyregistry.org/AB_1270780) |
| Rabbit polyclonal anti-SMARCC2 | Bethyl | Cat# A301-038A |
| Goat polyclonal anti-SOX2 | Santa Cruz | Cat# sc-17320  RRID: [AB_2286684](http://antibodyregistry.org/AB_2286684) |
| Rat monoclonal anti-SOX2 | eBioscience | Cat# 14-9811-82  RRID: AB_2924272 |
| Mouse monoclonal anti-TUBB3 | Covance | Cat# MMS-435P  RRID: [AB_2313773](http://antibodyregistry.org/AB_2313773) |
| Normal Rabbit IgG | Cell Signaling Technology | Cat# 2729  RRID: [AB_1031062](http://antibodyregistry.org/AB_1031062) |
| Normal Rabbit IgG | Cell Signaling Technology | Cat# 3900  RRID: [AB_1550038](http://antibodyregistry.org/AB_1550038) |
| Normal Mouse IgG | Cell Signaling Technology | Cat# 5415  RRID: [AB_10829607](http://antibodyregistry.org/AB_10829607) |
| Alexa Fluor donkey anti-rabbit IgG | Invitrogen | Cat# A21206 (488)  RRID: [AB_2535792](http://antibodyregistry.org/AB_2535792)  Cat# A21207 (594)  RRID: [AB_141637](http://antibodyregistry.org/AB_141637) |
| Alexa Fluor donkey anti-rabbit IgG | Jackson ImmunoResearch | Cat# 711-606-152  RRID: [AB_2340625](http://antibodyregistry.org/AB_2340625) |
| Alexa Fluor donkey anti-mouse IgG | Invitrogen | Cat# A21202 (488)  RRID: [AB_141607](http://antibodyregistry.org/AB_141607)  Cat# A21203 (594)  RRID: [AB_141633](http://antibodyregistry.org/AB_141633) |
| Alexa Fluor donkey anti-mouse IgG | Jackson ImmunoResearch | Cat# 715-606-151  RRID: [AB_2340866](http://antibodyregistry.org/AB_2340866) |
| Alexa Fluor donkey anti-goat IgG | Invitrogen | Cat# A21447 (647)  RRID: [AB_2535864](http://antibodyregistry.org/AB_2535864) |
| Alexa Fluor donkey anti-rat IgG | Invitrogen | Cat# A21208 (488)  RRID: [AB_2535794](http://antibodyregistry.org/AB_2535794)  Cat# A21209 (594)  RRID: [AB_2535795](http://antibodyregistry.org/AB_2535795) |
| Rabbit Anti-Mouse Immunoglobulins HRP | Dako | Cat# P0161  RRID: [AB_2687969](http://antibodyregistry.org/AB_2687969) |
| Goat Anti-Rabbit Immunoglobulins HRP | Dako | Cat# P0448  RRID: [AB_2617138](http://antibodyregistry.org/AB_2617138) |
| Anti-Rabbit IgG HRP | Rockland | Cat# 18-8816-31  RRID: [AB_2610847](http://antibodyregistry.org/AB_2610847) |
| Anti-Mouse Ig HRP | Rockland | Cat# 18-8817-31  RRID: [AB_2610850](http://antibodyregistry.org/AB_2610850) |
| **Biological samples** |  |  |
| Human fetal tissue | MRC-Wellcome Trust Human Developmental Biology Resource | http://hdbr.org |
| **Experimental models: Cell lines** | | |
| clone C1 of parental line HPSI0114ikolf2 | Wellcome Sanger Institute | [www.hipsci.org](http://www.hipsci.org) |
| KOLF2.1J | The Jackson Laboratory for Genomic Medicine | [www.hipsci.org](http://www.hipsci.org) |
| HPSI0214i-kucg_2 | Wellcome Sanger Institute | [www.hipsci.org](http://www.hipsci.org) |
| **Chemicals, peptides, and recombinant proteins** | | |
| Geltrex | ThermoFisher Scientific | Cat# A1413201 |
| Synthemax | Sigma-Aldrich | Cat# CLS3535 |
| DPBS | ThermoFisher Scientific | Cat# 14190-094 |
| DMEM/F-12 | ThermoFisher Scientific | Cat# 11320033 |
| Neurobasal Medium | ThermoFisher Scientific | Cat# 21103049 |
| N-2 Supplement | ThermoFisher Scientific | Cat# 17502001 |
| B-27 Supplement | ThermoFisher Scientific | Cat# 17504044 |
| GlutaMAX-l Supplement | ThermoFisher Scientific | Cat# 35050-038 |
| Penicillin/  Streptomycin | ThermoFisher Scientific | Cat# 15140122 |
| Y-27632 ROCK inhibitor | Tocris | Cat# 1254/10 |
| SB31542 | Abcam | Cat# ab120163 |
| LDN193189 | StemCell Technologies | Cat# 72147 |
| DAPT | Cambridge Bioscience | Cat# SM15-10 |
| Accutase | Sigma-Aldrich | Cat# A6964 |
| HBSS | ThermoFisher Scientific | Cat# 14170088 |
| tracrRNA | IDT | Cat# 1072533 |
| IDT duplex buffer | IDT | Cat# 11-05-01-12 |
| Alt-R® S.p. HiFi Cas9 Nuclease V3 | IDT | Cat# 1081060 |
| EP enhancer | IDT | Cat# 1075916 |
| CloneR | Stemcell Technologies | Cat# 05888 |
| Pierce IP lysis buffer | ThermoFisher Scientific | Cat# 87787 |
| Halt™ Protease Inhibitor Cocktail | ThermoFisher Scientific | Cat# 87786 |
| Halt™ Protease Inhibitor Cocktail | ThermoFisher Scientific | Cat# 78420 |
| BSA | ThermoFisher Scientific | Cat# 23209 |
| Sample Buffer, Laemmli 2× Concentrate | Sigma | Cat# S3401-10VL |
| 10x Tris/Glycine/SDS | Bio-Rad | Cat# 1610732 |
| 10x Tris Buffered Saline | Bio-Rad | Cat# 1706435 |
| Dried Skimmed Milk Powder | Marvel | N/A |
| ECL detection reagent | Amersham | Cat# RPN2236 |
| Protein G Sepharose^®^, Fast Flow | Sigma | Cat# P3296 |
| Paraformaldehyde, 4% in PBS | Alfa Aesar | Cat# J61899 |
| Live/Dead™ Fixable Near-IR Dead Cell Stain | Invitrogen | Cat# L34976 |
| Normal donkey serum | Jackson ImmunoResearch | Cat# 017-000-121 |
| DAPI | Sigma | Cat# D9564 |
| Vectashield Antifade Mounting Medium | Vector Laboratories | Cat# H-1000-10 |
| di(N-succimidyl) glutarate | Sigma-Aldrich | Cat# 80424 |
| Pierce™ 16% Formaldehyde (w/v), Methanol-free | ThermoFisher Scientific | Cat# 28908 |
| Glycine | Sigma-Aldrich | Cat# 50046 |
| Tris-HCl, pH 7.5 | In house^1^ | N/A |
| Tris, pH 8.0 | In house^1^ | N/A |
| EDTA | ThermoFisher Scientific | Cat# 87788 |
| EDTA | Sigma | Cat# E7889 |
| SDS | ThermoFisher Scientific | Cat# AM9820 |
| Triton X-100 | Sigma | Cat# T8787 |
| HEPES,  pH 7.9 | Sigma | Cat# H3375 |
| Sodium deoxycholate | Sigma | Cat# 30970 |
| LiCl | Sigma | Cat# L7026 |
| NP-40 Alternative | Sigma | Cat# 18896 |
| RNase A | ThermoFisher Scientific | Cat# E0531 |
| Proteinase K | ThermoFisher Scientific | Cat# 78437 |
| Tris-HCl, pH 7.4 | In house^1^ | N/A |
| MgCl_2_ | In house^1^ | N/A |
| NaCl | Sigma | Cat# S5150 |
| Water | ThermoFisher Scientific | Cat# AM9937 |
| **Critical commercial assays** | | |
| P3 Primary Cell 4D-Nucleofector^TM^ X Kit L | Lonza | Cat# V4XP-3024 |
| RNeasy Micro Kit | Qiagen | Cat# 74004 |
| Maxima First Strand cDNA Synthesis kit with dsDNase | ThermoFisher Scientific | Cat# K1671 |
| Taqman Universal qRT-PCR Master Mix | ThermoFisher Scientific | Cat# 4304437 |
| Pierce BCA Protein Assay Kit | ThermoFisher Scientific | Cat# 23225 |
| 4–15% Mini-PROTEAN® TGX™ Precast Protein Gels | Bio-Rad | Cat# 4561085 |
| Trans-Blot Turbo Mini 0.2 µm PVDF Transfer Packs | Bio-Rad | Cat# 1704156 |
| Hyperfilm™ ECL™ | Amersham | Cat# 28-9068-36 |
| Duoluk In Situ Red Started Kit Mouse/Rabbit | Sigma-Aldrich | Cat# DUO92101-1KT |
| KAPA mRNA polyA HyperPrep Kit | Illumina | Cat# KR1352 |
| Dynabeads Protein G | ThermoFisher Scientific | Cat# 10003D |
| Dynabeads Protein A | ThermoFisher Scientific | Cat# 10008D |
| KAPA pure beads | Roche | Cat# 07893271001 |
| Zymo Clean & Concetrator-5 Kit | Zymo Research | Cat# D4014 |
| NEB Ultra II DNA Library Prep Kit for Illumina | New England BioLabs | Cat# E7103 |
| Illumina Tagment DNA TDE1 Enzyme and Buffer Small Kit | Illumina | Cat# 20034197 |
| NEBNext HiFi 2X PCR Master Mix | New England BioLabs | Cat# M0541S |
| QubitTM dsDNA HS assay | ThermoFisher Scientific | Cat# Q32851 |
| **Oligonucleotides** | | |
| qRT-PCR Taqman Probes | ThermoFisher Scientific | Supplementary Table 2 |
| Primers for indexing ATAC-seq libraries | (Buenrostro et al., 2013) | Supplementary Table 3 |
| **Deposited data** | | |
| Raw and analyzed data | This paper | GEO: GSE214383 |
| Human reference genome NCBI build 37, GRCh37 | Genome Reference Consortium | http://www.ncbi.nlm.nih.gov/projects/genome/assembly/grc/human/ |
| **Software and algorithms** | | |
| All data analysis scripts used in this paper | This paper | [https://github.com/strohstern/op17_PM21134_ChIP.git](https://eur03.safelinks.protection.outlook.com/?url=https%3A%2F%2Fgithub.com%2Fstrohstern%2Fop17_PM21134_ChIP.git&data=05%7C01%7C%7C12349e8930c742041b1908daa227a6cc%7C4eed7807ebad415aa7a99170947f4eae%7C0%7C0%7C638000588419831601%7CUnknown%7CTWFpbGZsb3d8eyJWIjoiMC4wLjAwMDAiLCJQIjoiV2luMzIiLCJBTiI6Ik1haWwiLCJXVCI6Mn0%3D%7C3000%7C%7C%7C&sdata=6tH8azFDeC%2FGu7G2LYzyaZq8H8Vitz2StpH%2BuNKCXg4%3D&reserved=0)  [https://github.com/FrancisCrickInstitute/SC21030_OP_ASCL1ko](https://eur03.safelinks.protection.outlook.com/?url=https%3A%2F%2Fgithub.com%2FFrancisCrickInstitute%2FSC21030_OP_ASCL1ko&data=05%7C01%7C%7Cdb88460df0f3449e3de008daa625ccdb%7C4eed7807ebad415aa7a99170947f4eae%7C0%7C0%7C638004978530900385%7CUnknown%7CTWFpbGZsb3d8eyJWIjoiMC4wLjAwMDAiLCJQIjoiV2luMzIiLCJBTiI6Ik1haWwiLCJXVCI6Mn0%3D%7C3000%7C%7C%7C&sdata=zuh6uvSucT808eXIR8%2BS9awKBVSiRDlB2GeNd1yk3VY%3D&reserved=0) |
| Fiji v2.3.0/1.53q | (Schindelin et al., 2012) | <http://fiji.sc/> |
| FlowJo v10.8.1 | BD Life Sciences | [https://www.flowjo.com](https://www.flowjo.com/) |
| GraphPad Prism 9 v9.4.1 | GraphPad | <https://www.graphpad.com/> |
| Cutadapt v1.9.1 | (Martin, 2011) | <https://github.com/marcelm/cutadapt/> |
| RSEM v1.3.0 | (Li and Dewey, 2011) | <https://github.com/deweylab/RSEM> |
| STAR alignment algorithm v2.5.2a | (Dobin et al., 2013) | <https://github.com/alexdobin/STAR> |
| DESeq2 v1.12.3 | (Love et al., 2014) | <https://bioconductor.org/packages/release/bioc/html/DESeq2.html> |
| R | R Foundation | <https://www.r-project.org/> |
| DAVID Bioinformatics Resources | (Huang da et al., 2009; Sherman et al., 2022) | <https://david.ncifcrf.gov/summary.jsp> |
| nf-core/ChIP-seq pipeline v1.1.0 | (Ewels et al., 2020) | <https://doi.org/10.5281/zenodo.3529400> |
| nf-core/atacseq pipeline v1.1.0 | (Ewels et al., 2020) | <https://doi.org/10.5281/zenodo.3529420> |
| BEDTools v2.30.0 | (Quinlan and Hall, 2010) | <https://bedtools.readthedocs.io/en/latest/> |
| DeepTools v3.5.0 | (Ramírez et al., 2016) | <https://deeptools.readthedocs.io/en/develop/> |
| DiffBind v3.4.11 | (Ross-Innes et al., 2012) | <https://bioconductor.org/packages/release/bioc/html/DiffBind.html> |
| Activity-by-Contact algorithm | (Fulco et al., 2019) | [https://github.com/strohstern/op17_PM21134_ChIP.git](https://eur03.safelinks.protection.outlook.com/?url=https%3A%2F%2Fgithub.com%2Fstrohstern%2Fop17_PM21134_ChIP.git&data=05%7C01%7C%7C12349e8930c742041b1908daa227a6cc%7C4eed7807ebad415aa7a99170947f4eae%7C0%7C0%7C638000588419831601%7CUnknown%7CTWFpbGZsb3d8eyJWIjoiMC4wLjAwMDAiLCJQIjoiV2luMzIiLCJBTiI6Ik1haWwiLCJXVCI6Mn0%3D%7C3000%7C%7C%7C&sdata=6tH8azFDeC%2FGu7G2LYzyaZq8H8Vitz2StpH%2BuNKCXg4%3D&reserved=0) |
| Cell Ranger v5.0.0 | 10X Genomics | [https://www.10xgenomics.com](https://www.10xgenomics.com/) |
| Seurat v4.1.1 | (Butler et al., 2018; Hao et al., 2021; Stuart et al., 2019) | <https://satijalab.org/seurat/> |
| scVelo v0.2.2 | (Bergen et al., 2020) | <https://github.com/theislab/scvelo/blob/master/docs/source/index.rst> |
| HOMER v3.1 | (Heinz et al., 2010) | <http://homer.ucsd.edu/homer/motif/> |
| Interactive Genomics Viewer (IGV) v2.12.3 | (Robinson et al., 2011; Thorvaldsdottir et al., 2013) | <https://software.broadinstitute.org/software/igv/> |
| **Other** | | |
| Amaxa 4D Nucleofector | Lonza | Cat# AAF-1002X |
| Cell strainer-capped tubes | Falcon | Cat# 352235 |
| 1.5 ml Picoruptor Microtubes with Caps | Diagenode | Cat# C30010016 |
| Eppendorf^®^ LoBind microcentrifuge tubes | Fisher Scientific | Cat# 022431081 |
| Agilent TapeStation 4200 System | Agilent | Cat# G2991AA |
| Qubit 3.0 Fluorometer | Life Technologies | Cat# Q33216 |

^1^ = Media preparation STP, The Francis Crick Institute

**References to Resources Table**

Bergen, V., Lange, M., Peidli, S., Wolf, F.A., and Theis, F.J. (2020). Generalizing RNA velocity to transient cell states through dynamical modeling. Nat Biotechnol *38*, 1408-1414.

Buenrostro, J.D., Giresi, P.G., Zaba, L.C., Chang, H.Y., and Greenleaf, W.J. (2013). Transposition of native chromatin for fast and sensitive epigenomic profiling of open chromatin, DNA-binding proteins and nucleosome position. Nat Methods *10*, 1213-1218.

Butler, A., Hoffman, P., Smibert, P., Papalexi, E., and Satija, R. (2018). Integrating single-cell transcriptomic data across different conditions, technologies, and species. Nat Biotechnol *36*, 411-420.

Dobin, A., Davis, C.A., Schlesinger, F., Drenkow, J., Zaleski, C., Jha, S., Batut, P., Chaisson, M., and Gingeras, T.R. (2013). STAR: ultrafast universal RNA-seq aligner. Bioinformatics *29*, 15-21.

Ewels, P.A., Peltzer, A., Fillinger, S., Patel, H., Alneberg, J., Wilm, A., Garcia, M.U., Di Tommaso, P., and Nahnsen, S. (2020). The nf-core framework for community-curated bioinformatics pipelines. Nat Biotechnol *38*, 276-278.

Fulco, C.P., Nasser, J., Jones, T.R., Munson, G., Bergman, D.T., Subramanian, V., Grossman, S.R., Anyoha, R., Doughty, B.R., Patwardhan, T.A.*, et al.* (2019). Activity-by-contact model of enhancer–promoter regulation from thousands of CRISPR perturbations. Nature Genetics *51*.

Hao, Y., Hao, S., Andersen-Nissen, E., Mauck, W.M., 3rd, Zheng, S., Butler, A., Lee, M.J., Wilk, A.J., Darby, C., Zager, M.*, et al.* (2021). Integrated analysis of multimodal single-cell data. Cell *184*, 3573-3587 e3529.

Heinz, S., Benner, C., Spann, N., Bertolino, E., Lin, Y.C., Laslo, P., Cheng, J.X., Murre, C., Singh, H., and Glass, C.K. (2010). Simple combinations of lineage-determining transcription factors prime cis-regulatory elements required for macrophage and B cell identities. Mol Cell *38*, 576-589.

Huang da, W., Sherman, B.T., and Lempicki, R.A. (2009). Systematic and integrative analysis of large gene lists using DAVID bioinformatics resources. Nat Protoc *4*, 44-57.

Li, B., and Dewey, C.N. (2011). RSEM: accurate transcript quantification from RNA-Seq data with or without a reference genome. BMC Bioinformatics *12*.

Love, M.I., Huber, W., and Anders, S. (2014). Moderated estimation of fold change and dispersion for RNA-seq data with DESeq2. Genome Biology *15*.

Martin, M. (2011). Cutadapt removes adapter sequences from high-throughput sequencing reads. 2011 *17*, 3.

Quinlan, A.R., and Hall, I.M. (2010). BEDTools: a flexible suite of utilities for comparing genomic features. Bioinformatics *26*, 841-842.

Ramírez, F., Ryan, D.P., Grüning, B., Bhardwaj, V., Kilpert, F., Richter, A.S., Heyne, S., Dündar, F., and Manke, T. (2016). deepTools2: a next generation web server for deep-sequencing data analysis. Nucleic acids research *44*.

Robinson, J.T., Thorvaldsdottir, H., Winckler, W., Guttman, M., Lander, E.S., Getz, G., and Mesirov, J.P. (2011). Integrative genomics viewer. Nat Biotechnol *29*, 24-26.

Ross-Innes, C.S., Stark, R., Teschendorff, A.E., Holmes, K.A., Ali, H.R., Dunning, M.J., Brown, G.D., Gojis, O., Ellis, I.O., Green, A.R.*, et al.* (2012). Differential oestrogen receptor binding is associated with clinical outcome in breast cancer. Nature *481*, 389-393.

Schindelin, J., Arganda-Carreras, I., Frise, E., Kaynig, V., Longair, M., Pietzsch, T., Preibisch, S., Rueden, C., Saalfeld, S., Schmid, B.*, et al.* (2012). Fiji: an open-source platform for biological-image analysis. Nat Methods *9*, 676-682.

Sherman, B.T., Hao, M., Qiu, J., Jiao, X., Baseler, M.W., Lane, H.C., Imamichi, T., and Chang, W. (2022). DAVID: a web server for functional enrichment analysis and functional annotation of gene lists (2021 update). Nucleic Acids Res.

Stuart, T., Butler, A., Hoffman, P., Hafemeister, C., Papalexi, E., Mauck, W.M., 3rd, Hao, Y., Stoeckius, M., Smibert, P., and Satija, R. (2019). Comprehensive Integration of Single-Cell Data. Cell *177*, 1888-1902 e1821.

Thorvaldsdottir, H., Robinson, J.T., and Mesirov, J.P. (2013). Integrative Genomics Viewer (IGV): high-performance genomics data visualization and exploration. Brief Bioinform *14*, 178-192.

**Supplemental Table 4.** ATAC-seq primers with barcodes (Buenrostro et al., 2013).

| **Identifier** | **Sequence** |
| --- | --- |
| Ad1 noMX | AATGATACGGCGACCACCGAGATCTACACTCGTCGGCAGCGTCAGATGTG |
| Ad2.1 TAAGGCGA | CAAGCAGAAGACGGCATACGAGATTCGCCTTAGTCTCGTGGGCTCGGAGATGT |
| Ad2.2 CGTACTAG | CAAGCAGAAGACGGCATACGAGATCTAGTACGGTCTCGTGGGCTCGGAGATGT |
| Ad2.3 AGGCAGAA | CAAGCAGAAGACGGCATACGAGATTTCTGCCTGTCTCGTGGGCTCGGAGATGT |
| Ad2.4 TCCTGAGC | CAAGCAGAAGACGGCATACGAGATGCTCAGGAGTCTCGTGGGCTCGGAGATGT |
| Ad2.5 GGACTCCT | CAAGCAGAAGACGGCATACGAGATAGGAGTCCGTCTCGTGGGCTCGGAGATGT |
| Ad2.6 TAGGCATG | CAAGCAGAAGACGGCATACGAGATCATGCCTAGTCTCGTGGGCTCGGAGATGT |
| Ad2.7 CTCTCTAC | CAAGCAGAAGACGGCATACGAGATGTAGAGAGGTCTCGTGGGCTCGGAGATGT |
| Ad2.8 CAGAGAGG | CAAGCAGAAGACGGCATACGAGATCCTCTCTGGTCTCGTGGGCTCGGAGATGT |
| Ad2.9 GCTACGCT | CAAGCAGAAGACGGCATACGAGATAGCGTAGCGTCTCGTGGGCTCGGAGATGT |
| Ad2.10 CGAGGCTG | CAAGCAGAAGACGGCATACGAGATCAGCCTCGGTCTCGTGGGCTCGGAGATGT |
| Ad2.11 AAGAGGCA | CAAGCAGAAGACGGCATACGAGATTGCCTCTTGTCTCGTGGGCTCGGAGATGT |
| Ad2.12 GTAGAGGA | CAAGCAGAAGACGGCATACGAGATTCCTCTACGTCTCGTGGGCTCGGAGATGT |
| Ad2.13 GTCGTGAT | CAAGCAGAAGACGGCATACGAGATATCACGACGTCTCGTGGGCTCGGAGATGT |
| Ad2.14 ACCACTGT | CAAGCAGAAGACGGCATACGAGATACAGTGGTGTCTCGTGGGCTCGGAGATGT |
| Ad2.15 TGGATCTG | CAAGCAGAAGACGGCATACGAGATCAGATCCAGTCTCGTGGGCTCGGAGATGT |
| Ad2.16 CCGTTTGT | CAAGCAGAAGACGGCATACGAGATACAAACGGGTCTCGTGGGCTCGGAGATGT |
| Ad2.17 TGCTGGGT | CAAGCAGAAGACGGCATACGAGATACCCAGCAGTCTCGTGGGCTCGGAGATGT |
| Ad2.18 GAGGGGTT | CAAGCAGAAGACGGCATACGAGATAACCCCTCGTCTCGTGGGCTCGGAGATGT |

**Supplemental Table 5.** List of genes used to group DIV24 cells into three populations: cycling progenitors, transitional progenitors, and neurons.

| **Cycling Progenitors** | **Transitional Progenitors** | **Neurons** |
| --- | --- | --- |
| *PAX6*  *FABP7*  *VIM*  *NES*  *SOX2*  *MCM2*  *MKI67*  *TOP2A*  *UBE2C*  *SLC1A3*  *HES5*  *HES1* | *ASCL1*  *SOX4*  *DLX1*  *DLL1*  *RGS16*  *GADD45G*  *CDKN1C*  *HES6*  *SMOC1*  *DLL3*  *IGFBP5* | *TUBB3*  *DCX*  *ELAVL3*  *ELAVL4*  *MAP2*  *BCL11B*  *RBFOX3*  *NEFM*  *STMN1*  *STMN2* |
